# *Trsp* is required by regulatory T cells to prevent lethal autoimmunity in mice

**DOI:** 10.1101/2025.09.09.675163

**Authors:** Justin Jacobse, Jennifer M. Pilat, Anna Brooke Harris, Aaron Kwag, Zaryab Aziz, Channing Chi, Sam Schaefer, M. Diana Neely, Matthew A. Buendia, Andrew Pahnke, Christopher S. Williams, Wentao Deng, M. Kay Washington, Jeffrey C. Rathmell, Charles R. Flynn, Edmond H.H.M. Rings, Sarah P. Short, K. Sandeep Prabhu, Janneke N. Samsom, Jeremy A. Goettel, Yash A. Choksi

## Abstract

Selenoproteins are involved in immune cell metabolism, yet the roles of these proteins in T cell development and function remain largely unknown. The *Trsp* gene encodes the selenocysteine tRNA (tRNA^Sec^) required for translation of all selenoproteins. In this study, we found that *Trsp* was required for thymopoiesis, with the majority of tRNA^Sec^-deficient T cells not progressing beyond double negative 3 stage, with egressed thymocytes undergoing peripheral homeostatic expansion. *Trsp-*deficient CD4^+^ T cells exhibited impairments in TCR and IL-2 signaling and did not cause inflammation in experimental models. On the other hand, *Trsp*-deficient regulatory T (Treg) cells exhibited defects in suppressive function *ex vivo* and Treg-specific *Trsp* deletion using *Trsp*^fl/fl^*Foxp3*^YFP-Cre^ (*Trsp*^!ιTreg^) mice caused fatal autoimmunity similar to FOXP3-deficient mice. Reducing oxidative stress via 2-HOBA administration prolonged survival in these *Trsp*^!ιTreg^ mice. These findings indicate that tRNA^Sec^ is required for T cell homeostasis and may be therapeutic targets in inflammation.

**One sentence summary:** *Trsp*, a gene required for translation of all selenoproteins, is essential for all T cell development and function, especially regulatory T cells.

## Introduction

T cells are a critical component of adaptive immunity and their dysregulation is associated with various pathologies including infection, autoimmunity, and cancer. Both the development and function of T cells are tightly regulated by their metabolic capacity.(*1–6*) Specifically, degree of oxidative stress has been shown to regulate physiologic activity of T cells. For instance, low concentrations of reactive oxygen species (ROS) can promote T cell activation and proliferation, whereas higher ROS concentrations can inhibit transcription factors essential for T cell activation(*7*) and subsequent downregulation of transcription of interleukin-2.(*8*) One protein family that contributes to cellular metabolism and redox state is selenocysteine-containing proteins (selenoproteins), most of which are antioxidants.(*9*) Translation of the 25 known selenoproteins in humans (24 in mice) requires the selenocysteine tRNA (tRNA^Sec^) encoded by *TRU-TCA1-1*/*Trsp* along with specialized translation factors that collectively coordinate selenocysteine insertion.(*10*) Thus, *TRU-TCA1-1*/*Trsp* regulates the expression of the entire class of selenoproteins.(*11*) While human and murine T cells express selenoproteins, there is a preponderance for expression of specific family members, suggesting distinct cell-specific roles.(*12*)

Selenoproteins are essential for normal growth and development and although there are no reported null mutations in *TRU-TCA1-1*, individuals with a single nucleotide polymorphism (65C>G) that reduces tRNA^Sec^ levels and activity *in vitro* exhibit thyroid hormone imbalances.(*13*– *15*) In mice, deletion of *Trsp* is embryonic lethal.(*16*) Therefore targeted deletion of *Trsp* in specific cell types is required to investigate selenoprotein function collectively, which is particularly important as individual selenoproteins can compensate for each other.(*17*) In T cells, selenoproteins have previously been studied using proximal *Lck*^Cre^*-*mediated excision of *Trsp* that revealed a reduction in the frequency of mature CD4^+^/CD8^+^ single positive (SP) T cells in the thymus as well as in the periphery (here defined as extrathymic) and reduced activation/proliferation *ex vivo*.(*18*) While selenoproteins are known to be important for thymocyte development, it is currently unknown when selenoproteins are expressed during intrathymic T cell development and how this affects progression through each stage of thymopoiesis. Moreover, the metabolic requirements for T cells differ among various T cell subsets.(*19*) Although physiological levels of ROS are required for IL-2 production in activated T cells,(*20*, *21*) the regulatory roles of selenoproteins in T cell subtype function are not known. Thus, we sought to determine the effect of *en-bloc* deletion of murine selenoproteins in T cells on thymopoiesis and T cell function both *in vitro* and *in vivo*.

## Results

### CD2^+^ cells require *Trsp* for progression from DN3 to DN4 during thymopoiesis

To study the contributions of selenoproteins during thymopoiesis, mice deficient for *Trsp* in all T cells were generated by crossing *Trsp*^flox/flox^ mice(*22*) with *CD2*^iCre^ mice (*Trsp*^!ιCD2^ mice) to allow deletion of selenoproteins in double negative, CD4^+^CD8^+^, and CD4^+^ thymocytes. *Trsp*^flox/flox^ mice have loxP sites flanking the *Trsp* gene that encodes tRNA^Sec^ (*23*). Without tRNA^Sec^, selenoprotein transcripts are not translated.(*23*) To explore whether *Trsp-*deficiency affects T cell development, we examined the thymus in age-matched littermates. Compared to control mice, the thymus of *Trsp*^!ιCD2^ mice was nearly absent (**Figure 1A**), similar to what was reported in using *Lck*^Cre^*-* mediated excision of *Trsp*.(*24*) Correspondingly, the absolute number of thymocytes was decreased in *Trsp*^!ιCD2^ mice (**Figure 1B**). Thus, *Trsp* expression in CD2^+^ cells is required for thymopoiesis. To define which thymopoietic stages require *Trsp*, developing T cell subsets of control and *Trsp*^!ιCD2^ mice were defined by flow cytometry (FC). In the thymus, T cell development progresses through distinct stages before T cells become double positive (DP) and subsequently single positive (SP) CD4^+^ or CD8^+^ T cells (as reviewed in (*25*)). In *Trsp*^!ιCD2^ mice, the frequency of double negative (DN) thymocytes was increased with a concomitant decrease in the frequency of DP thymocytes (**Figure 1C**). This finding indicates that either the progression from DN to DP cells is dependent on *Trsp* expression, or that progression within the 4 distinct DN stages may be impaired when *Trsp* is absent in CD2^+^ cells, ultimately resulting in fewer DP cells. Lineage negative (CD19^neg^Ly6C^neg^TCRγο^neg^NK1.1^neg^CD11c^neg^) DN thymocyte stages are distinguished based on cell-surface expression of CD25 and CD44: DN1 (CD44^+^CD25^neg^), DN2 (CD44^+^CD25^+^), DN3 (CD44^low^CD25^+^), and DN4 (CD44^neg^CD25^neg^) as reviewed in (*26*). In *Trsp*^!ιCD2^ mice, the frequency of T cells at the DN3 stage was increased while cells at the DN4 stage were decreased compared to control mice (**Figure 1D**). Collectively, these data suggest that *Trsp* expression is required for progression through DN stages of thymocyte development with fewer cells progressing from DN3 to DN4 in *Trsp*^!ιCD2^ mice.

**Figure 1:**
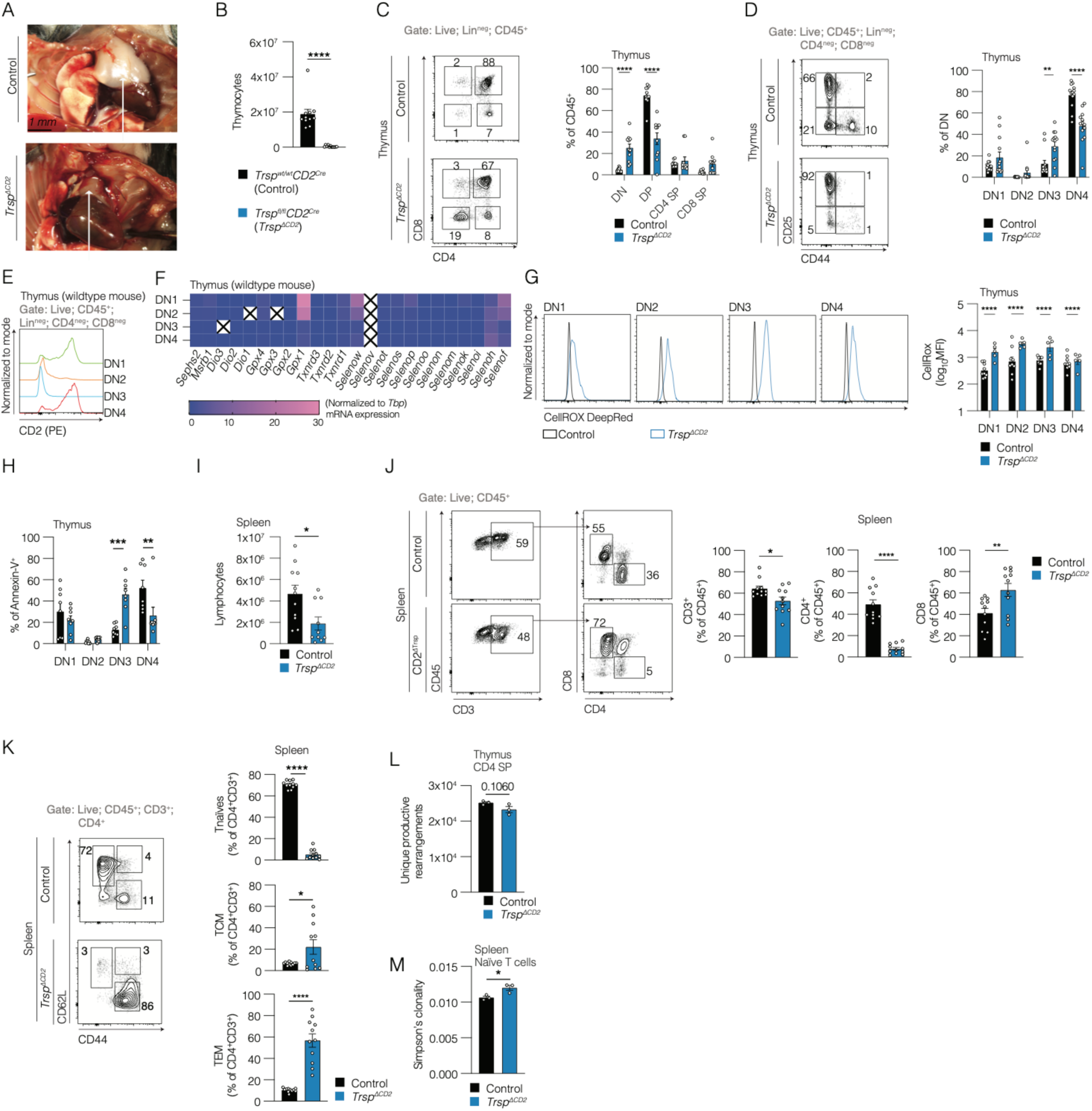
CD2^+^ cells require *Trsp* for thymopoiesis and T cell homeostasis. (A) Male *Trsp^fl/fl^* mice were crossed with female *Trsp^fl/wt^CD2^Cre^* mice to generate *Trsp^fl/fl^CD2^Cre^* (*Trsp^!ιCD2^*) mice. Representative photo of the thoracic cavity of a 5-week old control (*Trsp^wt^CD2^Cre^*) and *Trsp^!ιCD2^* mouse (age- and sex-matched). Arrows indicate thymus (black) and thymus remnant (blue). (B) Absolute cell count of live lymphocytes in the thymus acquired using automatic cell counter. Pooled data from 2 independent experiments. Unpaired t-test. (C) Representative FC plots (left) and quantification (right) of thymocytes. Pooled data from 2 independent experiments. Mixed model with post-hoc Šídák’s multiple comparisons test. Lineage^neg^ for thymocytes indicates the following markers: CD19^neg^Ly6C^neg^TCRγο^neg^NK1.1^neg^CD11c^neg^. (D) Representative FC plots (left) and quantification (right) of DN (CD4^neg^CD8^neg^) thymocyte subsets. Pooled data from 2 independent experiments. Mixed model with post-hoc Šídák’s multiple comparisons test. (E) FC histogram of CD2 expression in WT (*Trsp^wt^CD2^wt^*) mouse CD45^+^Lin^neg^CD4^neg^CD8^neg^ double negative thymocyte subsets. Representative results of 2 independent experiments. (F) Thymocytes were flow-sorted (DN1: CD45^+^CD4^neg^CD8^neg^CD44^+^CD25^neg^; DN2: CD45^+^CD4^neg^CD8^neg^CD44^+^CD25^+^; DN3: CD45^+^CD4^neg^CD8^neg^CD44^neg^CD25^+^; DN4: CD45^+^CD4^neg^CD8^neg^CD44^neg^CD25^neg^) from WT mice for selenoprotein RT-qPCR. Normalized expression is 2^-^*^!ι^*^C(t)^. Pooled data from 2 independent experiments with total n=6 biological replicates. (G) CellROX fluorescence in thymocyte subsets from control and *Trsp^!ιCD2^* mice. Representative histograms (left) and quantification (right). Data from 2 independent experiments performed using the same protocol including the same antibodies, same cytometer (Cytek), and same cytometer settings. Mixed model with post-hoc Šídák’s multiple comparisons test. (H) Annexin-V^+^ T cells gated and quantified into DN1-DN4 as in (D). Mixed model with post-hoc Šídák’s multiple comparisons test. (I) Absolute cell count of live lymphocytes in the spleen acquired using automatic cell counter. Pooled data from 2 independent experiments. Unpaired t-test. (J) Representative FC plots (left) and quantification (right) for CD4 and CD8 in splenocytes from control and *Trsp^!ιCD2^* mice. Pooled data from 2 independent experiments. Unpaired t tests. (K) Representative FC plots (left) and quantification (right) of splenic T cells (CD45^+^CD3^+^CD4^+^) from control and *Trsp^!ιCD2^* mice. Subsets defined as follows: TCM: CD44^+^CD62L^+^; Tnaïves: CD44^neg^CD62L^+^; TEM: CD44^+^CD62L^neg^. Pooled data from 2 independent experiments. Unpaired t tests. (L-M) Immunoseq was performed on flow-sorted (L) thymic CD4^+^ SP cells (CD45^+^CD4^+^CD8^neg^) and (M) splenic naïve T cells. Data from one experiment. Unpaired t-tests. * p<0.05; ** p<0.01; *** p<0.001; **** p<0.0001.

### CD2 and selenoproteins are expressed throughout thymopoiesis

To determine whether CD2 expression temporally coincides with alterations in the frequency of DN3 and DN4 populations in *Trsp*^!ιCD2^ mice, we examined *Cd2* expression across thymocyte stages in wild-type (WT) mice. We analyzed a previously published single-cell RNA-seq dataset of mouse DN thymocytes(*27*) and found that *Cd2* expression was readily detected in putative DN1 thymocytes (**Figure S01**). To confirm, CD2 expression was analyzed on thymic cells from WT mice. A bimodal distribution was observed, with CD2 being high in DN1, low/absent in DN2 and DN3, and high in DN4 (**Figure 1E**). This CD2 expression pattern is congruent with the concept of CD2 being required for signaling through the pre-T cell receptor (TCR) complex,(*28*) as DN3 is the first stage at which thymocytes start expressing TCR on their cell surface.(*29*)

Since CD2 is expressed during DN1, we profiled selenoprotein expression throughout thymopoiesis. To this end, we employed real-time quantitative reverse transcription polymerase chain reaction (RT-qPCR).(*30*) Several selenoproteins were expressed during all DN stages, including DN1. The selenoproteins with the highest expression levels were *Gpx1, Selenow, Selenoh,* and *Selenof*. Interestingly, certain selenoproteins such as *Gpx1*, *Selenow,* and *Selenof* were highly expressed in DN1/DN2 cells, whereas others (e.g., *Selenoh*) were only detected in later stages (**Figure 1F**). To examine whether human thymocytes exhibited a similar expression pattern, we analyzed a single-cell RNA-seq dataset of human thymocytes(*31*) and observed that some of the same selenoproteins were highly expressed in earlier DN stages (*GPX1*, *SELENOH, SELENOF*) compared with later stages (**Figure S02**). Thus, select selenoproteins are expressed by developing thymocytes in mice and humans and may be required for proper thymocyte development.

### Thymocytes require *Trsp* to maintain physiological redox balance and prevent aberrant apoptosis

Having determined that *Trsp*^!ιCD2^ mice exhibit altered thymopoiesis, we next interrogated the potential mechanism underlying this observation. Most selenoproteins function by reducing oxidative stress, which can induce apoptosis (reviewed in (*32*)). Thymocyte apoptosis is central to thymopoiesis as T cells that do not receive a permissive TCR signal die instead of developing into DP cells. We hypothesized that selenoproteins function to reduce oxidative stress-induced apoptosis during thymopoiesis. First we measured ROS in *Trsp*^!ιCD2^ mice by FC using CellROX. As expected, ROS levels were higher during all DN stages in *Trsp*^!ιCD2^ mice as compared to control mice (**Figure 1G**). Since the thymi in *Trsp*^!ιCD2^ mice were smaller than in WT controls, we hypothesized that this increase in ROS may drive aberrant apoptosis. As CellROX histograms did not segregate into distinct negative and positive populations (**Figure 1G**), we were not able to compare the frequency of Annexin-V^+^ cells in cells with high versus low levels of ROS. Therefore, we examined which DN cells undergo apoptosis by measuring the frequency of Annexin-V^+^ cells. In WT mice, the majority of Annexin-V^+^ cells were in the DN4 stage, whereas in *Trsp*^7CD2^ mice most of the Annexin-V^+^ cells were observed at the DN3 stage (**Figure 1H**), consistent with the tRNA^Sec^-deficient T cells not progressing beyond DN3. Collectively, these data suggest that CD2^+^ cells require selenoproteins to maintain physiological redox state for appropriate thymocyte development through the DN stages.

### tRNA^Sec^-deficient T cells that egress from the thymus of *Trsp***^!ι^**^CD2^ mice undergo homeostatic proliferation

Having established that *Trsp*^!ιCD2^ mice have decreased thymic output, we wanted to determine if this leads to lymphopenia. We examined the spleen and mesenteric lymph nodes (mLN) of *Trsp*^!ιCD2^ mice and found a reduction in the absolute number of lymphocytes in the spleen of *Trsp*^!ιCD2^ mice with lower frequencies of CD4^+^ T cells and higher frequencies of CD8^+^ T cells compared to control mice (**Figure 1I-J**). We then examined CD4^+^ T cell subsets, (i.e., naïve T (TN), T effector memory (TEM) or T central memory (TCM) cells) and found that while TN frequency was reduced, both TEM and TCM frequencies were increased (**Figure 1K**). Similar findings were observed in the mLNs, with the notable exceptions that there was no change in CD3^+^ T cells, and there were no increases in CD8^+^ T or TCM cell frequencies (**Figure S03**). This pattern of altered CD4^+^ T cell subsets with increased memory T cells and decreased naïve T cells is consistent with peripheral tRNA^Sec^-deficient CD4^+^ T cells undergoing homeostatic proliferation.(*33*) Furthermore, the TCR repertoire during thymic development and post-egress determined via TCR CDR3 sequencing revealed that the number of productive TCRs in the CD4^+^ SP thymocytes trended lower in tRNA^Sec^-deficient T cells (**Figure 1L**) while the clonality of the splenic naïve T cells was increased (**Figure 1M**). This is consistent with a TCR repertoire of *Trsp*^!ιCD2^ mice that is less diverse than WT mice. These data indicate that selenoprotein-deficient T cells fail to undergo appropriate positive selection in the thymus and that the few T cells that successfully egress undergo homeostatic proliferation in the periphery.

### Pathogenic CD4^+^ T cells require *Trsp*

Our observations indicate that T cells require *Trsp* for proper T cell development. Since *Trsp*^!ιCD2^ mice have dysregulated CD4^+^/CD8^+^ T cells, we generated mice that allow for inducible deletion of selenoproteins specifically in CD4^+^ T cells (*Cd4*^Cre-ERT2^*;Rosa26*^lox-stop-lox-tdTomato^*;Trsp*^fl/fl^ mice). Repeated tamoxifen gavage resulted in >90% CD4^+^ T cells expressing tdTomato, indicative of Cre-mediated recombination (**Figure S04A**). We confirmed the downstream consequences of *Trsp* deletion by Western blot for the selenoproteins glutathione peroxidase 4 (GPX4) and thioredoxin reductase 1 (TXNRD1), and observed marked reductions in GPX4 and TXNRD1 protein levels in *Trsp*^!ιCD4^ as compared to WT CD4^+^ T cells (**Figure S04B**). To test whether T cells require *Trsp* for pathogenicity, we employed the adoptive transfer model of colitis(*34*) wherein naïve T cells (live, CD4^+^CD45RB^high^) from *Trsp*^!ιCD4^ or control mice are injected into lymphocyte-deficient *Rag1^-/-^* recipient mice (**Figure 2A**). The frequency of injected tdTomato^+^ cells did not differ between tRNA^Sec^-deficient or -sufficient naïve T cells (**Figure 2B**), indicating that a similar frequency of cells underwent inducible Cre-mediated excision of the *Rosa26* lox-stop-lox sequence prior to adoptive transfer. At the experimental endpoint, *Rag1*^-/-^ mice that received tRNA^Sec^-deficient naïve T cells were protected against colitis (**Figure 2C**). This was not attributed to differences in engraftment or expansion, as the frequency of splenic CD4^+^CD3^+^ T cells did not differ between recipient groups (**Figure 2D**). Moreover, tRNA^Sec^-deficient T cells recovered from recipient mice and restimulated *ex vivo* with PMA/ionomycin produced IFNγ to a similar degree as tRNA^Sec^- sufficient T cells (**Figure 2E**). Since T cell activation is a critical driver of colitis, this finding suggests that *Trsp* deficiency may lead to a TCR signaling defect, as cytokine production remained intact when bypassing TCR signaling with PMA/ionomycin. To test this, we utilized a T cell-independent colitis model mediated by dextran-sodium sulphate (DSS) and found no differences in weight loss, colon length, or histological injury between *Trsp*^!ιCD4^ and control mice (**Figures 2F-H**). Thus, *Trsp* is required for CD4^+^ T cell pathogenicity *in vivo*, likely downstream of TCR activation.

**Figure 2:**
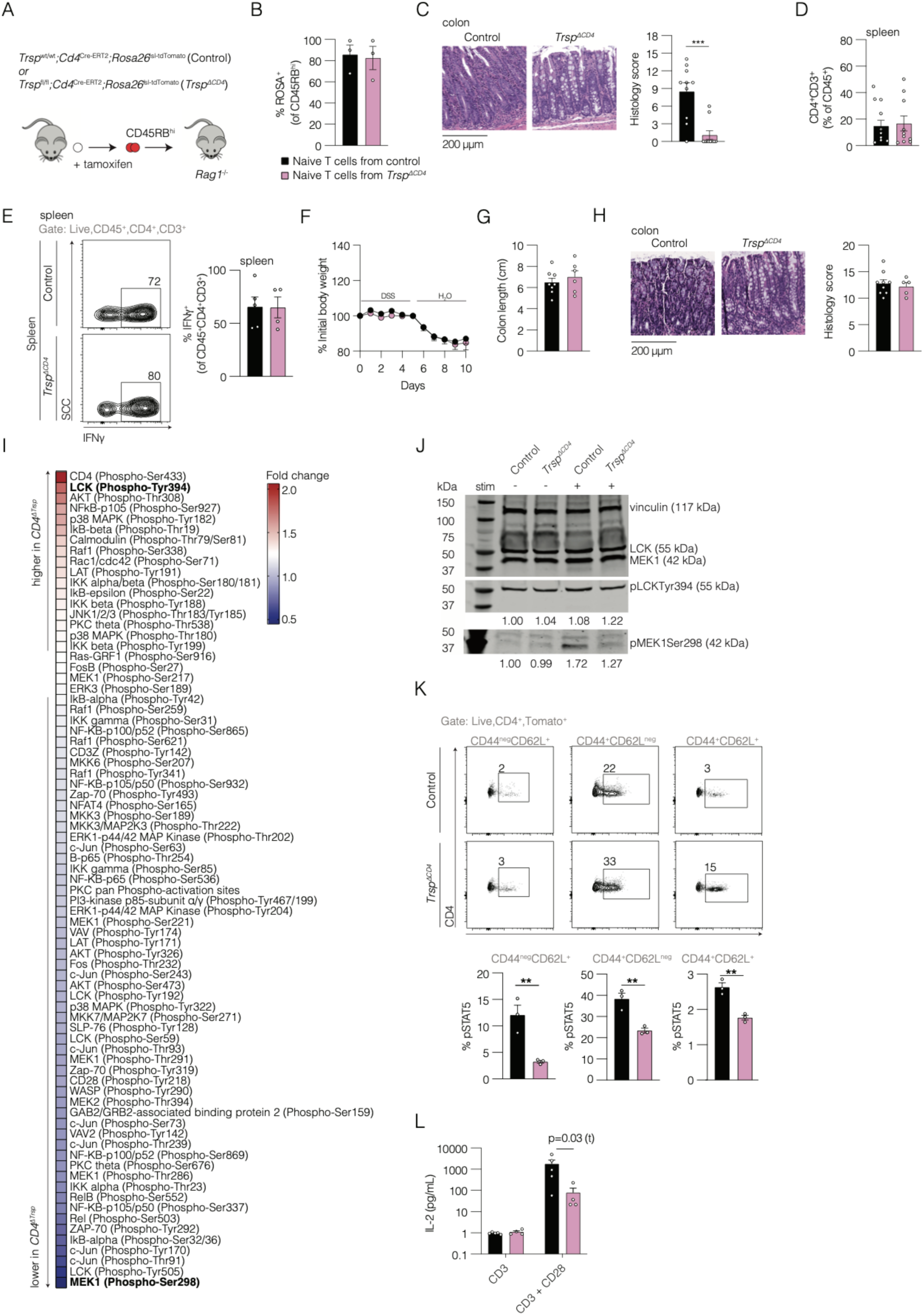
T cells require *Trsp* for TCR activation and pathogenicity. (A) *_Cd4_Cre-ERT2_;ROSA_lox-stop-lox-tdTomato_;Trsp_wt/wt* _(control) and *Cd4*_*Cre-ERT2_;ROSA_lox-stop-lox-tdTomato_;Trsp_fl/fl* _(*Trsp*_*!ιCD4*_)_ mice were gavaged with 8 mg tamoxifen on days 0, 1 and 3. Naïve T cells (CD45RB^high^) were then flow-sorted and intraperitoneally injected into sex-matched *Rag1^-/-^* mice. (B) Frequency of ROSA^+^ T cells in sorted cells prior to injection into *Rag1^-/-^* recipients. Pooled data from 3 independent experiments. Within each experiment, samples were pooled per genotype prior to sorting; each datapoint represents 3-6 mice. Mann-Whitney U test. (C) Representative histology (left) and quantification (right) of colitis across 3 independent experiments 7 weeks after injection of naïve T cells. Mann-Whitney U test. (D) Frequency of splenic T cells in mice that received *Trsp*-deficient or -sufficient naïve T cells. Measured at experimental endpoint. Mann-Whitney U test. (E) Representative FC plots (left) and quantification (right) of splenic T cells restimulated with PMA/ionomycin for 5 hours in the presence of Golgi-Stop and stained for IFNγ. Mann-Whitney U test. (F) Baseline-normalized weight curves for control and *Trsp^!ιCD4^* mice gavaged with tamoxifen (8 mg x 3 doses) that subsequently received 3.5% DSS in their drinking water for 5 days followed by 5 days regular water, after which mice were euthanized. Mann-Whitney U test. (G) Colon length at experimental endpoint. Mann-Whitney U test. (H) Representative histology (left) and colitis score (right). Mann-Whitney U test. (I) CD4^+^ T cells were magnetically enriched from spleen and mLN of control and *Trsp^!ιCD4^* mice and stimulated using plate-bound αCD3 and soluble αCD28 for 2 hours, then lysed and subjected to a T cell phosphoprotein assay. Shown are the fold changes comparing *Trsp*-deficient to -sufficient T cells. n=9 per genotype; one experiment. (J) Western blot of control and *Trsp^!ιCD4^* CD4^+^ T cells stimulated as in (I) for 15 minutes for LCK, MEK1, pLCK^Tyr394^, pMEK1^Ser298^, and vinculin (loading control). Representative data from 2 independent experiments. (K) Representative FC plots (top) and quantification (bottom) of control and *Trsp^!ιCD4^* CD4^+^ T cells stimulated with IL-2 for 15 minutes and stained for pSTAT5. Subsets defined as follows: TCM: CD44^+^CD62L^+^; Tnaives: CD44^neg^CD62L^+^; TEM: CD44^+^CD62L^neg^. Representative data from 2 independent experiments. Mann-Whitney U tests. (L) ELISA for IL-2 on supernatant of control and *Trsp^!ιCD4^* CD4^+^ T cells after 2 days in culture. Unpaired t test. * p<0.05; ** p<0.01; *** p<0.001; **** p<0.0001.

### *Trsp* is required for T cell receptor activation

To determine experimentally whether *Trsp* modulates pathways downstream of TCR activation, *Trsp*-sufficient/deficient CD4^+^ T cells were stimulated using αCD3 and αCD28 followed by measurement of 213 TCR-relevant antibodies using a phosphoprotein assay. When ranked by fold change, phosphorylation of MEK1 at Ser298 (decreased) and phosphorylation of LCK at Tyr394 (increased) were identified as the two most altered in tRNA^Sec^-deficient as compared to control CD4^+^ T cells (**Figure 2I**). These results were corroborated via Western blot analysis (**Figure 2J**). In addition, IL-2 production is downstream from TCR activation, and IL-2 is an important regulator of T cell metabolic function.(*35*) Thus, we assessed whether *Trsp* deletion in CD4^+^ T cells alters both response to IL-2 and production of IL-2 after activation. Stimulation with exogenous IL-2 *in vitro* resulted in reduced phosphorylation of signal transducer and activator of transcription 5 (pSTAT5) (**Figure 2K**). Interestingly, not only was the response to exogenous IL-2 reduced, but IL-2 production was also reduced in *Trsp*^!ιCD4^ as compared to control CD4^+^ T cells following activation (**Figure 2L**). Altogether, these data show that CD4^+^ T cells require *Trsp* for activation of several signaling pathways downstream of TCR engagement. Moreover, tRNA^Sec^- deficient CD4^+^ T cells secrete less IL-2 after TCR activation and are less responsive to exogenous IL-2. These findings likely explain the inability of *Trsp*-deficient naïve CD4 T cells to cause colitis in *Rag1*^-/-^ mice.

### tRNA^Sec^-deficient Treg cells exhibit an activated phenotype but are functionally impaired Since

*Trsp* was required for TCR activation in *Trsp*^!ιCD4^ mice, and *Trsp*^!ιCD2^ mice were lymphopenic, we next sought to identify other CD4^+^ T cell subsets that require *Trsp*. We hypothesized that the function of *Trsp* in Treg cells differs from other CD4^+^ T cell subsets as we observed an increase in the frequency of splenic FOXP3^+^ Treg cells in *Trsp*^!ιCD2^ mice (**Figure 3A**). *Trsp*^!ιCD2^ Treg cells also exhibited a more active and proliferative phenotype based on expression of MHCII, Ki67, and CD69 (**Figure 3B**). Expression of RORγt was also increased in both CD4^+^FOXP3^+^ (**Figure 3B**) and CD4^+^FOXP3^neg^ T cells from *Trsp*^!ιCD2^ mice (**Figure S05**). Interestingly, CD4^+^FOXP3^neg^, but not CD4^+^FOXP3^+^, cells from *Trsp*^!ιCD2^ mice also expressed elevated levels of CD25, the high-affinity component of the IL-2R, which may lead to higher affinity for IL-2 as compared to their WT counterparts (**Figure S05**). This may, in part, explain the increase in proliferation of CD4^+^FOXP3^neg^ T cells of *Trsp*^!ιCD2^ mice as measured by Ki67 (**Figure S05**). Prior work established that RORγt^+^ Treg cells are more suppressive then RORγt^neg^ Treg cells(*36*, *37*). Therefore, we assessed the suppressive capacity of Treg cells from *Trsp*^!ιCD2^ mice *in vitro* and found that suppression by tRNA^Sec^-deficient Treg cells was modestly impaired compared to WT Treg cells (**Figure 3C**). Collectively, these data suggest that *Trsp* plays cell-intrinsic roles in Treg cell function.

**Figure 3:**
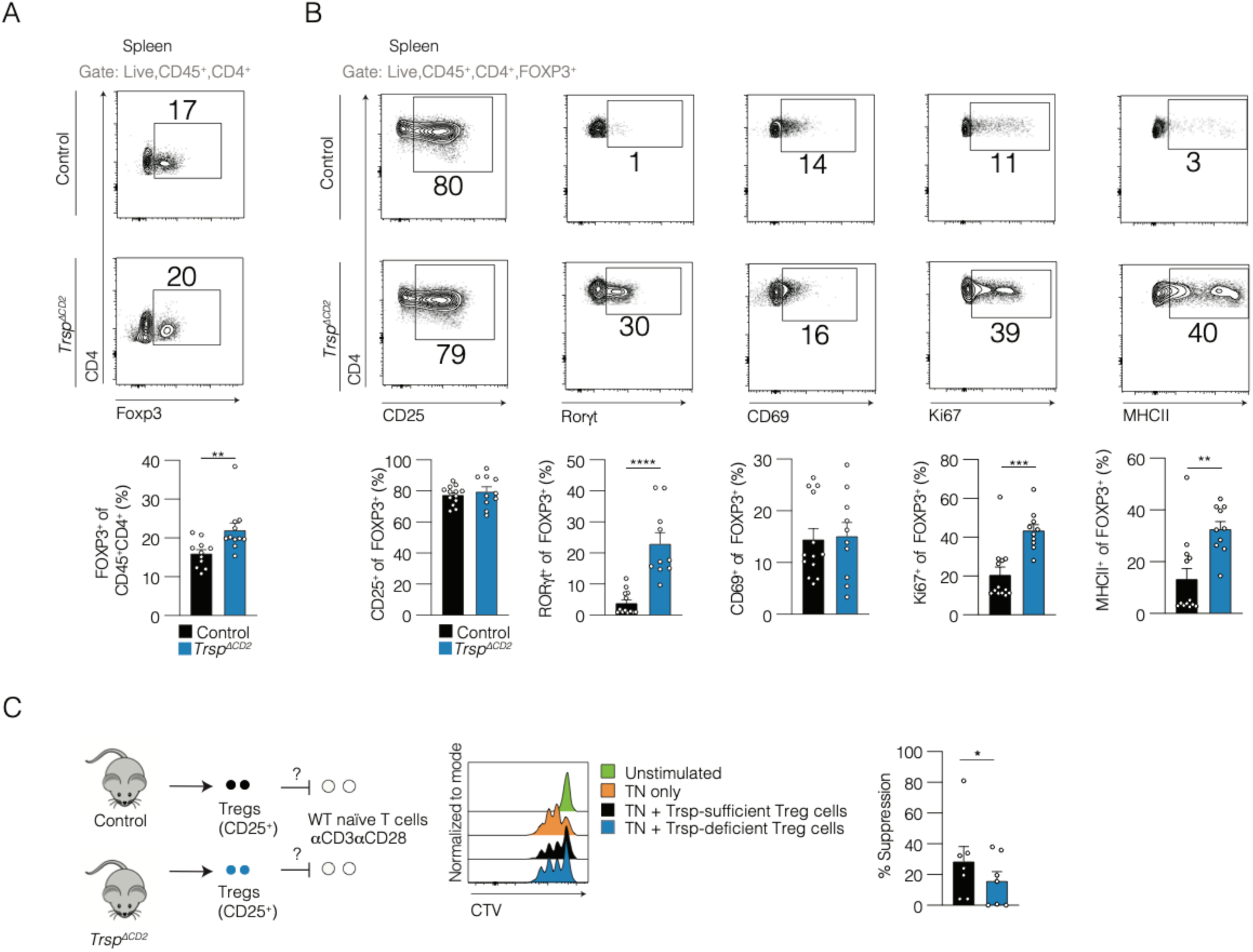
Treg cells in *Trsp^!ιCD2^* mice have an activated phenotype and reduced suppression. (A) Representative FC plots (top) and quantification (bottom) of splenic T cells from control and *Trsp^!ιCD2^* mice. Pooled data from 2 independent experiments. Unpaired t-test. (B) Representative FC gating plots (top) and quantification (bottom) of splenic Treg cells from control and *Trsp^!ιCD2^* mice. Pooled data from 2 independent experiments. Unpaired t tests. (C) Schematic (left), representative FC histograms (middle), and quantification (right) of *in vitro* Treg cell suppression using flow-sorted CD25^+^ T cells from control and *Trsp^!ιCD2^* mice and naïve T cells from control mice in a 1:1 ratio of Treg: naïve T cells. Each pair of dots (i.e. control and *Trsp^!ιCD2^*-derived Treg cells) represents one individual experiment. One-sided paired t test. * p<0.05; ** p<0.01; *** p<0.001; **** p<0.0001.

### Mice with tRNA^Sec^-deficient Treg cells develop fatal autoimmunity

Although *Trsp*^!ιCD2^ mice exhibit impaired thymopoiesis, they do not spontaneously develop overt inflammation. Nevertheless, they do exhibit increased Treg cell frequency, and these Treg cells appear to be activated with increased RORγt^+^ expression, yet they display defects in suppressive function *in vitro*. To investigate these Treg cell functional defects directly, we generated *Trsp*^fl/fl^*Foxp3*^YFP-Cre^ (*Trsp*^!ιTreg^) mice. Homozygous deletion of *Trsp* specifically in FOXP3^+^ cells resulted in an autoimmune phenotype (i.e. scaly tail, early mortality, skin inflammation, and hair loss) similar to FOXP3-deficient *Foxp3*^scurfy^ mice (**Figure 4A-B**).(*38*) Of note, gross phenotypes of *Trsp*^!ιTreg^ mice differed slightly from those of *Foxp3^scurfy^* mice, in that *Trsp*^!ιTreg^ mice did not develop the characteristic thickened ears (not shown). *Trsp*^!ιTreg^ mice also displayed thymic involution and multiorgan inflammation, as observed in *Foxp3^scurfy^* mice (**Figure 4C-D**), along with splenomegaly, decreased lymphocytes, and decreased splenic Treg cells (**Figure 4E-F**). Importantly, this reduction in splenic Treg cells could not be attributed to increased apoptosis, as measured by Annexin-V staining (**Figure S06**). Additionally, *Trsp*^!ιTreg^ and control mice displayed similar frequencies of splenic CD3^+^, CD8^+^, and CD4^+^ cells (**Figure 4F**). This systemic inflammation was also associated with decreased splenic B cells and increased serum IgE levels (**Figure 4G**), both characteristics of autoimmune inflammation, and similar to *Foxp3*^scurfy^ mice (**Figure S07**). Of note, *Trsp*^!ιTreg^ mice did not develop spontaneous colitis, which is also not reported in *Foxp3^scurfy^*mice(*39*).

**Figure 4:**
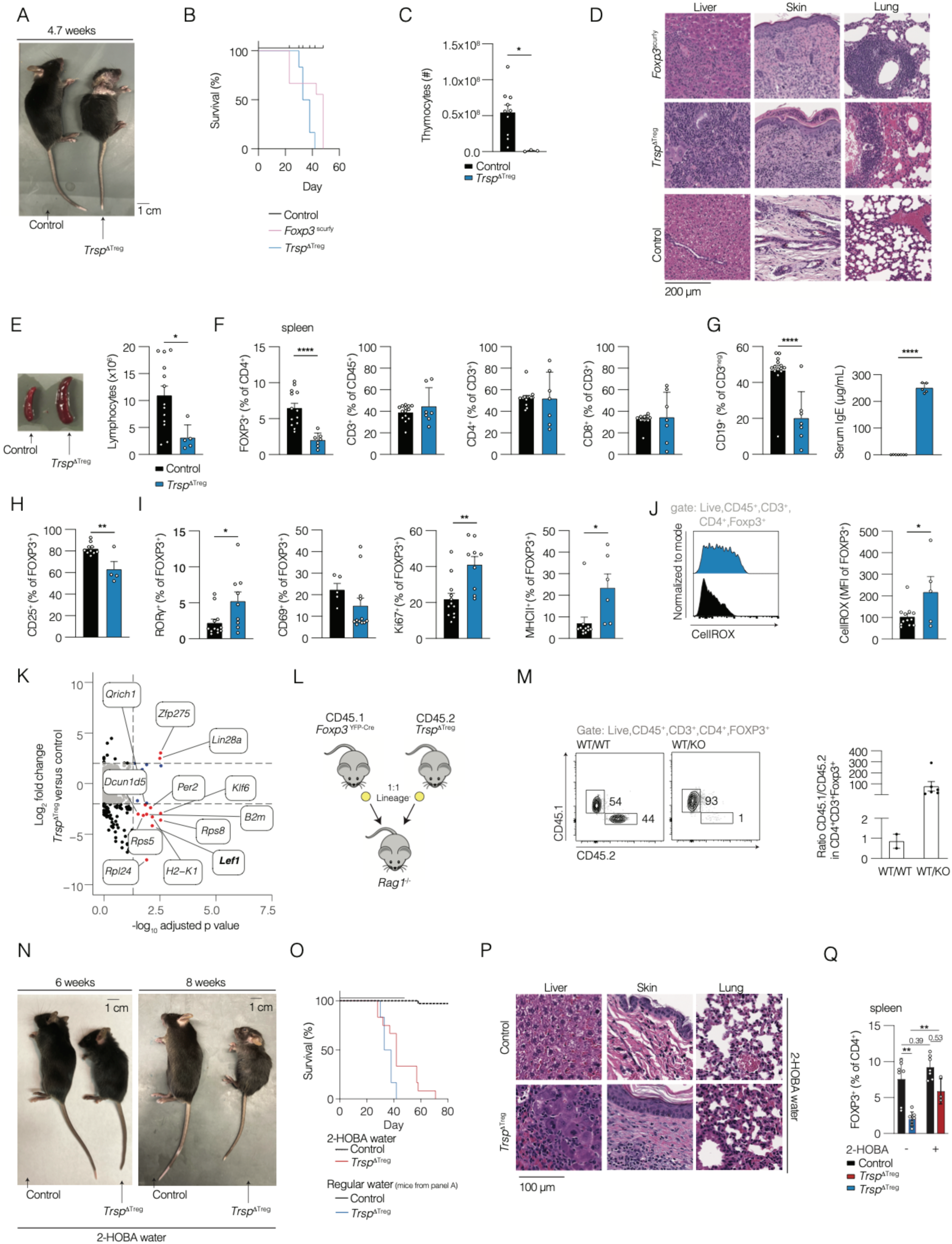
Treg cells require *Trsp* to prevent lethal autoimmunity. *Trsp^fl/fl^Foxp3^YFP-Cre^* mice were generated by crossing *Foxp3^YFP-Cre^* mice with *Trsp^fl/fl^* mice. (A) Representative photo of control and *Trsp*^fl/f*l*^*Foxp3*^YFP-Cre^ mouse at 4.7 weeks of age. (B) Survival curve of control, *Trsp^fl/fl^Foxp3^YFP-Cre^*, and *Foxp3^scurfy^* mice with Kaplan-Meier analysis. Pooled data from 5 independent experiments. n=22 control, n=9 *Foxp3^scurfy^,* n=6 *Trsp^fl/fl^Foxp3^YFP-Cre^.* Log-Rank (Mantel-Cox) test, p<0.0001. (C) Absolute cell count of live thymocytes from control and *Trsp^fl/fl^Foxp3^YFP-Cre^* mice acquired using automatic cell counter at 4 weeks of age. Pooled data from 3 independent experiments. Unpaired t-test. (D) Representative histology of liver, skin and lung of control, *Trsp^fl/fl^Foxp3^YFP-Cre^*, and *Foxp3^scurfy^* mice. (E) Photo of spleen (left) and absolute cell count (right) of live splenic lymphocytes from control, *Trsp^fl/fl^Foxp3^YFP-Cre^*, and *Foxp3^scurfy^* mice. Representative data from 3 independent experiments. (F-J) FC phenotyping of (F) splenic T cells, (G) splenic B cells and serum IgE, and (H-J) splenic Treg cells. Pooled data from 4 independent experiments. Unpaired t-tests. (K) RNA-sequencing was performed on FC-sorted splenic Treg cells from control and *Trsp^fl/fl^Foxp3^YFP-Cre^*mice. DESEQ2 was used to assess differentially expressed genes comparing *Trsp^fl/fl^Foxp3^YFP-Cre^* versus control Treg cells. n=3 control and n=*4 Trsp^fl/fl^Foxp3^YFP-Cre^*. (L) Schematic representing bone marrow chimera experiment. *Rag1^-/-^* mice were sublethally irradiated (450 rads) and transplanted using a 1:1 mixture of lineage-depleted bone marrow cells from *Foxp3^YFP-Cr^*^e^CD45.1 and *Trsp^fl/fl^Foxp3^YFP-Cre^CD45.2* mice. Control mice received a 1:1 mixture of cells from *Foxp3^YFP-Cre^CD45.1* and *Foxp3^YFP-Cre^CD45.2* mice. (M) Recipient mice were analyzed 6 weeks post-transplantation with the YFP^+^ fraction gated and the CD45.2^+^/CD45.1^+^ ratio quantified. Data representative of 2 independent experiments with *N=3-6*. (N-Q) *Trsp^fl/fl^Foxp3^YFP-Cre^* and control mice received 2-HOBA in the drinking water starting at postnatal day 1-2. (N) Representative photo of control and *Trsp^fl/fl^Foxp3^YFP-Cre^* mice at indicated ages. (O) Survival curve of control and *Trsp^fl/fl^Foxp3^YFP-Cre^*mice treated without (same data from B) or with 2-HOBA from birth with Kaplan-Meier analysis. Log-rank (Mantel-Cox) test. n=*22* control without 2-HOBA, n=*6 Foxp3^YFP-Cre;^ Trsp^fl/fl^* without 2-HOBA, n=62 control with 2-HOBA, n=12 *Foxp3^YFP-Cre;^ Trsp^fl/fl^*with 2-HOBA p<0.0001. (P) Representative histology of liver, skin and lung of control and *Foxp3^YFP-Cre;^ Trsp^fl/fl^* mice euthanized 58 days after 2-HOBA treatment. (Q) FC quantification of splenic Treg cells from control and *Trsp^fl/fl^Foxp3^YFP-Cre^* mice treated without (same data from F) or with 2-HOBA from birth at the end of experiment. Mixed model with post-hoc Šídák’s multiple comparisons test. p<0.05; ** p<0.01; *** p<0.001; **** p<0.0001.

CD4^+^FOXP3^+^ Treg, but not CD4^+^FOXP3^neg^, cells from *Trsp*^!ιTreg^ mice also exhibited an activated and proliferative phenotype (i.e. increased Ki67 and MHCII), along with increased ROS as measured via CellRox (**Figures 4I-J, S08**). To ascertain transcriptomic differences between *Trsp*-sufficient/deficient cells, we performed RNA-seq on sorted YFP^+^ (FOXP3^+^) cells from *Trsp*^!ιTreg^ and control mice. One gene that was significantly downregulated in tRNA^Sec^-deficient Treg cells was the WNT transcription factor *Lef1* (**Figure 4K**)*. Lef1* is both a WNT transcription factor and a WNT target gene and absence of *Lef1* renders Treg cells unable to suppress.(*40*) The decreased expression of *Lef1* in tRNA^Sec^-deficient Treg cells was confirmed by RT-qPCR (**Figure S09**). Since the frequency of Treg cells in *Trsp*^!ιTreg^ mice was reduced as compared to control mice (**Figure 4F**), we investigated whether *Trsp* expression provides a competitive advantage to Treg cells. Using congenically-marked mixed bone marrow chimeras, we found that the majority of Treg cells that persist in conditioned *Rag1^-/-^* mice were *Trsp*-sufficient (**Figures 4L-M**).

Given elevated oxidative stress in Treg cells of *Trsp*^!ιTreg^ mice, we tested whether the autoimmune phenotypes could be rescued via antioxidant treatment. During ROS-induced lipid peroxidation, toxic isoleuvoglandins are formed that can be scavenged by 2-hydroxybenzylamine (2-HOBA), reducing ROS-induced damage.(*41*) 2-HOBA is currently under clinical investigation for several autoimmune/inflammatory disorders.(*42*) Thus, we tested whether 2-HOBA in the drinking water could ameliorate the autoimmune phenotypes of *Trsp*^!ιTreg^ mice. At 6 weeks, *Trsp*^!ιTreg^ mice that received 2-HOBA were still smaller in size compared to *Trsp*-sufficient mice but did not develop skin ulcerations, lethargy, or scaly tail (**Figure 4N)** that were observed in *Trsp*^!ιTreg^ mice without 2-HOBA treatment by 5 weeks of age (**Figure 4A**). However, by 8 weeks *Trsp*^!ιTreg^ mice receiving 2-HOBA did begin to develop these phenotypes. Survival of *Trsp*^!ιTreg^ mice receiving 2-HOBA was prolonged compared to untreated *Trsp*^!ιTreg^ mice (**Figure 4O**) and inflammation in the skin and lung was less severe in 2-HOBA-treated *Trsp*^!ιTreg^ mice compared to untreated (**Figure 4P**). Splenic Treg cell numbers in *Trsp*^!ιTreg^ mice were higher with 2-HOBA treatment compared to *Trsp*^!ιTreg^ mice not administered 2-HOBA (**Figure 4Q**). Importantly, 2-HOBA treatment had no effect on splenic Treg numbers in control mice (**Figure 4Q**). Altogether, these data indicate that Treg expression of selenoproteins is required to prevent fatal autoimmunity in mice, possibly attributed to impaired *Lef1* expression and increased oxidative stress.

### Treg cells require *Trsp* for stability, suppression, and survival in an IL-2-dependent manner

Treg cells can lose *Foxp3* expression and consequently their suppressive function to become pathogenic ex-Treg cells.(*43*, *44*) It is possible that the systemic inflammation observed in *Trsp*^!ιTreg^ mice is due to conversion of tRNA^Sec^-deficient Treg cells to pathogenic ex-Tregs. In female *Trsp*^fl/fl^*Foxp3*^YFP-Cre/WT^ mice, which have both *Trsp*-sufficient and -deficient Treg cells due to random X-inactivation(*45*), the majority of CD4^+^CD25^+^ T cells were YFP^neg^ (∼75%), as compared to *Foxp3*^YFP-Cre/WT^ control mice, in which 50% of CD4^+^CD25^+^ T cells were YFP^neg^ (data not shown). These data suggest that Treg cells lose their identity in *Trsp*^fl/fl^*Foxp3*^YFP-Cre^ mice. To examine how loss of *Trsp* affects Treg cells and their propensity towards becoming ex-Treg cells, we generated lineage-tracing *Trsp*^fl/fl^*Foxp3*^eGFP-Cre-ERT2^*Rosa26*^lox-stop-lox-tdTomato^ reporter mice and compared them to control *Foxp3*^eGFP-Cre-ERT2^;*Rosa26*^lox-stop-lox-tdTomato^ mice. In these mice, FOXP3-expressing cells are constitutively marked by GFP expression, and, upon tamoxifen administration, become irreversibly labeled with tdTomato. Repeated tamoxifen treatment reduced splenic Treg cell frequency in *Trsp*^fl/fl^*Foxp3*^eGFP-Cre-ERT2^*Rosa26*^lox-stop-lox-tdTomato^ as compared to control mice (**Figures 5A-B**). The frequency of ex-Treg cells (tdTomato^+^GFP^neg^) trended upwards in *Trsp*^fl/fl^*Foxp3*^eGFP-Cre-ERT2^*Rosa26*^lox-stop-lox-tdTomato^ mice, but did not achieve statistical significance (**Figure 5C**). When we tested GFP^+^tdTomato^+^ Treg cells for suppressive function, we again observed that tRNA^Sec^-deficient Treg cells had a trend towards reduced suppressive capacity (**Figure 5D**). We then determined if loss of Treg cell stability (i.e. conversion to ex-Tregs) associated with *Trsp* deficiency is sufficient to cause inflammation in lymphopenic mice. To directly test this, GFP^+^tdTomato^+^ Treg cells were sorted and transferred into *Rag1^-/-^* recipient mice (**Figure 5E**). Both *Trsp*-deficient and -sufficient Treg cells did not cause weight loss or colitis (**Figure 5F** and data not shown), indicating that tRNA^Sec^-deficient (ex-)Treg cells are not pathogenic in a cell-intrinsic manner. However, when compared to mice that received (WT) naïve T cells, the frequency of Treg cells in mice that received tRNA^Sec^-sufficient Treg cells was increased to a greater extent as compared to mice that received tRNA^Sec^-deficient Treg cells (**Figure 5G**), suggesting that in a non-competitive setting, Treg cells need *Trsp* for sustenance.

**Figure 5:**
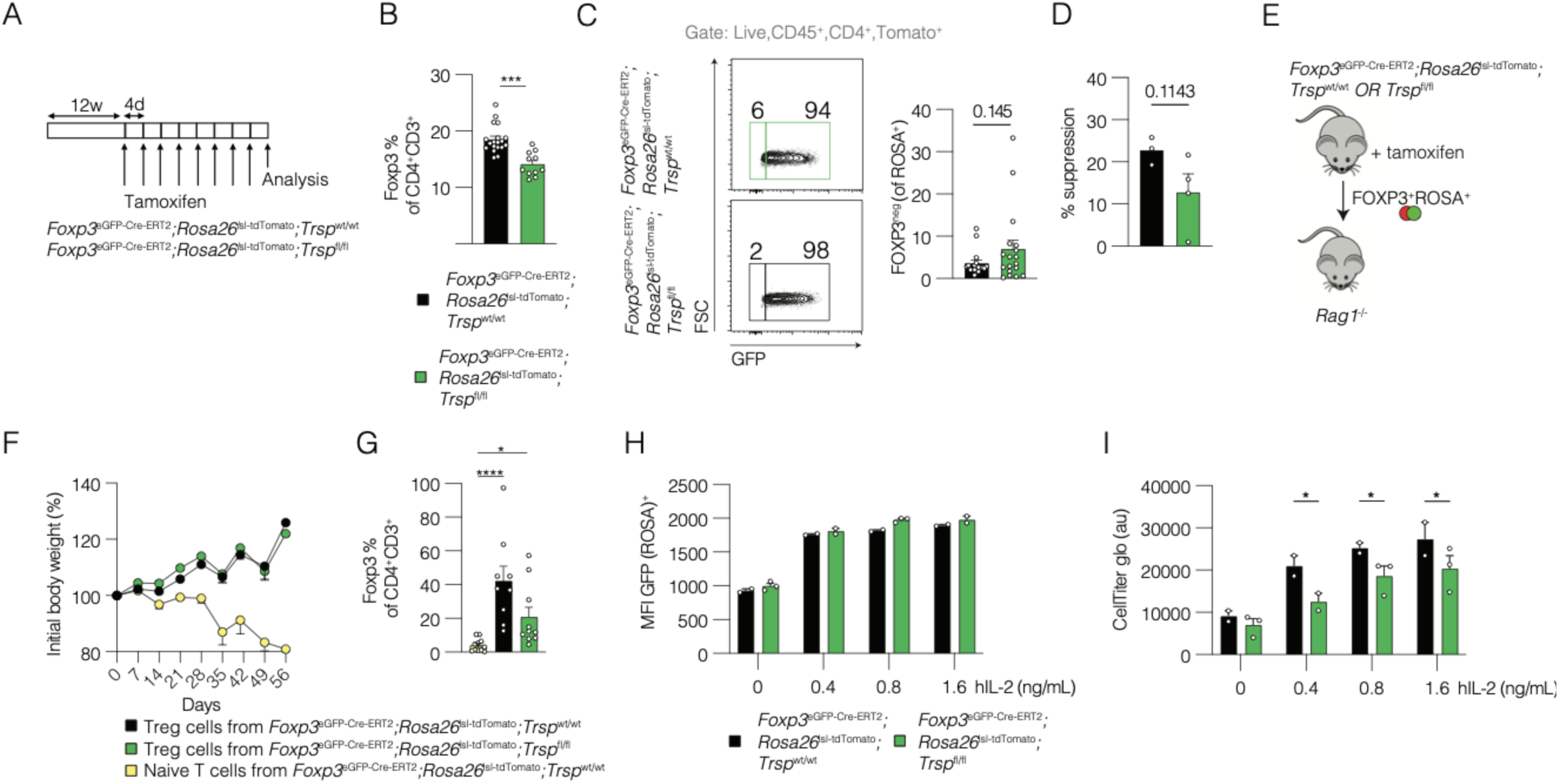
Treg cells require *Trsp* for stability, suppression, and survival in an IL-2-dependent manner. (A) *Foxp3^eGFP-Cre-ERT2;^ROSA^lox-stop-lox-tdTomato;^Trsp^wt/wt^* (control) and Foxp3*^eGFP-Cre-ERT2^*;*ROSA*^lox-stop-lox-tdTomato^;*Trsp^fl/fl^*were gavaged with 8 mg tamoxifen 8 times 4 days apart. (B) Splenic Treg cell frequency in mice from (A). Unpaired t-test. (C) Representative FC plots of ex-Treg cells (left) and quantification (right). Pooled data from 3 independent experiments. Unpaired t-test. (D) *In vitro* Treg cell suppression using flow-sorted GFP^+^tdTomato^+^ Treg cells from control and *Foxp3^eGFP-Cre-ERT2^;ROSA^lox-stop-lox-tdTomato^;Trsp^fl/fl^*mice and naïve T cells from control mice in a 1:1 ratio of Treg:naïve T cells. Pooled data from 2 independent experiments. Unpaired t-test. (E-G) Treg cells (ROSA^+^GFP^+^) were adoptively transferred into *Rag1^-/-^* mice. These mice did not develop colitis (not pictured). (E) Schematic showing experimental design. (F) Weight curve of *Rag1^-/-^* mice that received control naïve T cells, control Treg cells, or *Trsp^ΔTreg^* Treg cells. n=7-8 per condition. (G) Splenic Treg cell frequency in *Rag1^-/-^* mice that received control naïve T, control Treg cells, or *Trsp^ΔTreg^* Treg cells at the experimental endpoint. Kruskal-Wallis test. (H-I) Splenic Treg cells from control and *Foxp3^eGFP-Cre-ERT2^Rosa26 ^lox-stop-lox-tdTomato^; Trsp^fl/fl^* mice were FC sorted and cultured *in vitro* with indicated concentrations of human recombinant IL-2 for 48 hours before (H) FC. One experiment. Mixed model with post-hoc Šídák’s multiple comparisons test. (I) CellTiter glow quantification of cells in (H). Data was normalized to media only control. Representative data from two independent experiments. Mixed model with post-hoc Šídák’s multiple comparisons test. * p<0.05; ** p<0.01; *** p<0.001; **** p<0.0001.

As Treg cells are uniquely sensitive to IL-2 due to expression of the trimeric, high-affinity IL-2R, our observations in *Trsp*^7Treg^ mice raised the question of whether the advantage of *Trsp*- sufficient Treg cells is IL-2-dependent. To this end, splenic Treg cells were sorted from *Trsp*^fl/fl^*Foxp3*^eGFP-Cre-ERT2^*Rosa26*^lox-stop-lox-tdTomato^ mice and *Foxp3*^eGFP-Cre-ERT2^*Rosa26*^lox-stop-lox-tdTomato^ mice and cultured in the presence of increasing doses of IL-2 without TCR stimulation. No differences were observed in the MFI of GFP (FOXP3) in tdTomato^+^ tRNA^Sec^-deficient *versus* - sufficient Treg cells (**Figure 5H**), indicating that *Trsp* is not required for IL-2-mediated FOXP3 stability *in vitro*. Nevertheless, survival of tRNA^Sec^-deficient Treg cells was reduced as compared to tRNA^Sec^ -sufficient Treg cells in the presence of varying concentrations of IL-2 (**Figure 5I**). In summation, these data show that *Trsp* is required for Treg cell suppression, Treg cell stability (IL-2-independent), and Treg cell survival (IL-2-dependent).

## Discussion

In this report, we demonstrate that *Trsp* is required for normal thymopoiesis, peripheral CD4^+^ T cell activation, and regulatory T cell survival and function. More specifically, we illustrate that *Trsp* deletion in CD2^+^ cells impairs thymopoiesis by reducing progression of T cell progenitors from DN3 to DN4, resulting in lymphopenia and T cell exhaustion. In peripheral CD4^+^ T cells, *Trsp* deficiency impairs TCR activation, IL-2 signaling, and effector T cell function. Finally, we show that *Trsp* deletion specifically in Treg cells causes Treg cell deficiency and fatal autoimmune inflammation through increased oxidative stress. These phenotypes recapitulate those of mice with loss-of-function mutations in *Foxp3*.(*46–48*) As *Trsp* encodes a tRNA required for translation of all selenoproteins, these data position *Trsp* and selenoproteins as central to T cell physiology.

An earlier study suggested that *Trsp* expression is required for *Lck*-expressing thymocytes, yet did not define the consequences of *Trsp* deletion on thymopoeisis.(*18*) We found that *Trsp* is particularly important in the DN3 to DN4 transition and that selenoproteins (particularly *Gpx1, Selenow,* and *Selenof*) are expressed as early as DN1, while expression of *Txnrd1* in (all) DN thymocytes is relatively low as compared to other selenoproteins. Others have shown that WT non-*Cre*-expressing DN1 thymocytes have a competitive advantage over *Cre*-expressing *Txnrd1-*deficient DN1 thymocytes after transplantation into WT irradiated mice.(*49*) Despite increased oxidative stress in DN1 thymocytes, we did not observe this reduced DN1 cellularity in *Trsp*^fl/fl^*CD2*^Cre^ mice. Though these are seemingly discrepant findings, the advantage of WT DN1 thymocytes as compared with *Txnrd1-*deficient DN1 thymocytes was measured in a different context (i.e. post-irradiation and transfer as compared with from birth). Nevertheless, our data is consistent with their finding that selenoproteins may be relevant for pre-TCR signaling, as we demonstrated that there is CD2 expression in murine DN thymocytes and found it is highly expressed by DN1 cells, thus supporting a role for CD2 pre-TCR, as has been suggested.(*28*) Overall, there is sparse literature on how individual selenoproteins affect thymopoeisis, and much more work is required to determine their roles.

The observation that *Trsp*^!ιCD2^ mice have substantial aberrations in thymopoiesis and peripheral T cell proportions, studying the functional consequences of *Trsp* deletion in conventional CD4^+^ T cells was only possible utilzing inducible *Cre* based approaches. For example, memory T cells respond differently to external stimuli, i.e. TCR stimulation, then naïve T cells, and *Trsp*^!ιCD2^ mice have significantly more memory T cells than WT mice. Thus, to test the function of *Trsp* in conventional CD4^+^ T cells, we utilized cells isolated from *Cd4*^Cre-ERT2^*;Rosa26*^lox-stop-lox-tdTomato^*;Trsp*^fl/fl^ mice. To mimic TCR stimulation *in vitro*, we treated these *Trsp*-sufficient and -deficient CD4^+^ T cells using αCD3/αCD28, since T cells require signal via the CD3 protein complex and costimulatory signal via the CD28 protein complex, which is typically provided by antigen-presenting cells. We found that phosphorylation of MEK1 on Serine 298 is decreased in *Trsp*-deficient CD4^+^ T cells, which suggests decreased MEK1 autophosphorylation, decreased ERK activity,(*50*) and thus decreased T cell function.(*51*) This finding is consistent with a prior publication which found that ERK signaling was reduced in *Trsp-*deficient T cells.(*18*) We also found that phosphorylation of LCK on Tyrosine 394 is increased in *Trsp*-deficient CD4^+^ T cells, whereas phosphorylation of LCK on Tyrosine 505 is decreased, suggesting that LCK activity is increased in the absence of *Trsp*.(*52*) This is in accordance with others who have shown that T cells require *Selenok* for calcium signaling downstream of TCR signaling.(*53*) To what extent *Trsp-*deficient CD4^+^ T cell function can be rescued by restoring individual components of the TCR signaling cascade remains to be determined.

Alterations in TCR signaling lead to functional impairment as the adoptive transfer of *Trsp*-deficient naïve T cells into *Rag1^-/-^* mice did not cause colitis, likely due to disabling TCR-dependent activation. Consistent with this conclusion, cytokine production itself is intact when TCR is bypassed in these adoptively transferred cells. Yet, *Trsp* function in CD4^+^ T cells does not seem limited to TCR activation *per se*. *Trsp*-deficient CD4^+^ T cells respond to IL-2 stimulation with less phosphorylation of STAT5 then *Trsp*-sufficient CD4^+^ T cells. This observation is true across all CD4^+^ T cell subsets, including in antigen-inexperienced naïve T cells and in the absence of TCR activation. We did not test whether established colitis in the adoptive transfer model can be (partially) rescued by *Trsp* deletion, but such an experiment would be important to begin to explore whether modulation of *Trsp* in CD4^+^ T cells might be a therapeutic option to reduce inflammation in patients with autoimmune diseases.

In immune homeostasis, Treg cells suppress conventional CD4^+^ T cells. Similar to tRNA^Sec^-deficient conventional T cells, tRNA^Sec^-deficient Treg cells were also dysfunctional and lead to fatal autoimmunity. This finding is significant, as only deletion of CTLA-4(*54*) and IL-2R(*55*) subunits have previously been shown to phenocopy the autoimmunity observed in *Foxp3^scurfy^* mice. Understanding the mechanisms behind tRNA^Sec^-deficient Treg cell dysfunction is crucial. Our data suggest that increased oxidative stress renders Treg cells incapable of effective suppression, since 2-HOBA partially rescues this phenotype. *In vitro* suppression data indicate that Treg cells require *Trsp* during TCR-mediated activation and further investigation into how these processes occur is ongoing. Our findings seem to oppose reports that GPX1-deficient Treg cells are superior in suppressing T cell proliferation *in vitro*.(*56*) Yet, tRNA^Sec^-deficient Treg cells may be dissimilar from GPX1-deficient Treg cells in several aspects such as levels of oxidative stress and expression of other compensatory selenoproteins. Similar to conventional CD4^+^ T cells, tRNA^Sec^-deficient Treg cells also exhibit impaired responses to IL-2. IL-2 signaling is particularly important for Treg cell function, as they require IL-2(*55*) produced by activated effector T cells. *Ex vivo*, *Trsp-*deficient Treg cells demonstrate decreased survival as compared to *Trsp-*sufficient Treg cells. Based on these observations, *Trsp* may be a target for reducing the immune suppressive function of Treg cells in cancers, potentially allowing for increased immunosurveillance.

Important next steps in the field will be to determine which selenoproteins are required for differentiation into specific T cell subsets.(*57*) The extent to which specific selenoproteins affect individual Th subsets and Treg cell function have not been fully defined. While it has been shown that *Gpx1^-/-^* mice displayed enhanced Th1/Th17 polarization with a concomitant decrease in Th2 polarization,(*58*) and that *Gpx1^-/-^* CD25^+^ Treg cells) suppressed naïve T cell proliferation induced by αCD3 and WT dendritic cells to a greater degree than WT CD25^+^ Treg cells *in vitro*,(*56*) there is little data systematically determining which selenoproteins are most important in each Th subset. These are areas of ongoing investigation in our laboratory.

### Limitations of study

One limitation of this study is the deletion of all selenoproteins by targeting *Trsp*. Although this approach prevents confounding from compensatory effects of other selenoproteins, it also hinders the ability to distinguish the contributions of individual selenoproteins. Another limitation is that this study largely focused on the role of selenoproteins in the murine immune system and the translational relevance of these findings remains to be determined. Finally, for part of our studies we used a *CD2*^Cre^ mouse. CD2 expression is not limited to T cell progenitors, as it is also expressed by B cells. This is likely important as others have implicated selenoproteins in B cell development.(*59*, *60*)

## Material and Methods

### Mice

Mice lacking *Trsp* in specific immune cells were generated by crossing *Trsp*^flox/flox^ (*22*) mice with various *Cre* lines: *CD2^Cre^*mice(*61*) (The Jackson Laboratory, Bar Harbor, ME, JAX stock #008520); *Cd4*^Cre-ERT2^ mice(*62*) (JAX stock #022356); *Foxp3*^eGFP-Cre-ERT2^ mice(*63*) (JAX stock #016961); and *Foxp3*^YFP-Cre^ mice(*64*) (JAX stock #016959). As *Foxp3* is located on the X chromosome, we used *Foxp3^YFP-Cre^* hemizygous males and homozygous females for all experiments. To enable lineage tracing, the tamoxifen-inducible *Cre* mice (i.e. *Cd4*^Cre-ERT2^ and *Foxp3*^eGFP-Cre-ERT2^) were crossed with *Rosa26*^lox-stop-lox-tdTomato^ mice(*65*) (JAX stock #007914). Mice were administered 8 mg tamoxifen (Millipore Sigma, St. Louis, MO, #T5648) in 200 μL olive oil (Millipore Sigma, O1514) with 1% ethanol (Millipore Sigma, #1085430250) by oral gavage. To prevent bias from tamoxifen gavage and its associated *Cre* recombination(*66*), both knockout and control mice received tamoxifen. *Trsp*^flox/flox^*Foxp3*^YFP-Cre^ (and control) mice were monitored by independent observers blinded to experiment and genotype, and euthanized when deemed moribund. Macroscopic photos were taken using an iPhone (Apple, Cupertino, CA).

Genotyping for *Trsp* was done in-house using the following primers (Millipore Sigma) and the following protocol: CKNO2: 5’-GCAACGGCAGGTGTCGCTCTGCG-3’; 8RP: 5’- CGTGCTCTCTCCACTGGCTCA-3’. PCR conditions: 94°C for 5 min; 30 cycles of: 94°C for 30 sec, 58°C for 60 sec, 72°C for 60 sec; 72°C for 3 min. Using these primers and protocol, the floxed *Trsp* gene yields a ∼1.1 kb fragment, while the WT *Trsp* gene yields a 900 bp fragment. The Cre-excised *Trsp* gene yields a ∼450 bp fragment. Confirmatory genotyping was done by Transnetyx using their proprietary probes after we confirmed these probes yielded the same results as our in-house genotyping.

*Foxp3^scurfy^* (JAX stock #004088)(*67*) mice have been described previously. All mice except *Rag1*^- /-^ mice were housed in the same room. *Rag1*^-/-^ mice(*68*) (JAX stock #002216) were housed in sterilized cages and provided autoclaved food and water. All experiments were performed using 6–8-week-old sex- and age-matched, co-housed mice unless otherwise indicated. Both female and male mice were used. Littermates were used whenever possible. Experiments were approved by the VA Institutional Animal Care and Use Committee (IACUC) and/or the VUMC IACUC.

### Flow Cytometry

#### Staining

For cell surface staining, cells were blocked using mouse Fcblock (Biolegend, San Diego, CA, #101320) and incubated in the antibody cocktail including a fixable live/dead stain (ThermoFisher Scientific, Waltham, MA, #65-0866-14) for 20 minutes at 4°C in the dark. Red blood cells were lysed prior to staining where applicable using ACK lysing buffer (ThermoFisher Scientific, #A1049201). Intracellular cytokine staining was performed using Cytofix/Cytoperm (BD, #554714) and intranuclear stain was performed using the Foxp3/Transcription Factor Staining Buffer Set (ThermoFisher Scientific, #00-5523-00), the Foxp3/Transcription Factor staining buffer kit (Tonbo/Cytek, #TNB-0607-KIT), or the TrueNuclear^TM^ Transcription Factor Buffer Set (#424401, Biolegend), according to manufacturer’s instructions. Cells were stained for oxidative stress using CellROX Deep Red (ThermoFisher Scientific, #C10422) for 30 minutes at 37°C with 5% CO_2_ in 200 μL T cell media (TCM, RPMI 1640 with 10% heat-inactivated FBS, 100 unitsmL^-1^ penicillin, 100 μgmL^-1^ streptomycin (ThermoFisher Scientific, #10378016), GlutaMax (ThermoFisher Scientific, #35050-061), MEM non-essential amino acids (Corning, #25-025-CI), 55 μM beta-mercaptoethanol (ThermoFisher Scientific, #21985-023), and 1 mM sodium pyruvate (Millipore Sigma, #S8636-100ML). Stock vials of CellROX were used a maximum of two times as our observations showed similar results (not shown). Cells were stained for Annexin-V for 15 minutes at room temperature in either 1X Annexin-V binding buffer for Annexin-V antibody staining or PBS for Apo-15 peptide staining (Biolegend). Antibodies used for flow cytometry or cell sorting are listed in **Table S1**.

#### IL-2 stimulation and pSTAT5 staining

To measure T cell activation in response to IL-2, CD4^+^ T cells were enriched from spleen and mLN using immunomagnetic negative selection (EasySep™ Mouse CD4+ T Cell Isolation Kit, STEMCELL Technologies, #19852). T cells were stained for surface markers (**Table S1**) as above, washed with TCM, incubated for 5 minutes at 37°C, 5% CO2, then stimulated with 20 ngmL^-1^ recombinant mouse IL-2 (R&D Systems, #202-IL-010/CF) for 15 minutes at 37° C, 5% CO2. After stimulation, cells were fixed with warm Phosflow^TM^ Fix Buffer I (BD, # 557870) for 10 minutes at 37°C. After washing, cells were permeabilized with ice-cold (−20°C) Phosflow^TM^ Perm Buffer III (BD, # 558050) for 30 minutes on ice. Cells were washed again and stained for pSTAT5 for 1 hour at room temperature, then washed and acquired.

#### Acquisition and sorting

Flow cytometric analysis was performed using a 4/5-Laser Fortessa or 5-laser LSRII (BD, Franklin Lakes, NJ) with FACSDiva software (BD), or a Cytek® Aurora (CytekBio, Bethesda, MD). FC-sorting was performed on a Cytek Aurora^TM^ CS System or on a FACS Aria III (BD).

#### Adoptive transfer

For adoptive transfer colitis, naïve T cells were sorted from spleen and mLN based on live CD4^+^CD45RB^hi^ expression following pre-enrichment for CD4^+^ cells as above. *Rag1^-/-^* mice were intraperitoneally injected with 5x10^5^ naïve T cells.(*34*) As colitis in the adoptive transfer model is dependent on a complex microbiota containing *Helicobacter spp*., recipients were removed from sterile housing conditions and bedding from other (non-experimental) mice housed in non-autoclaved cages was mixed into the cages of recipient mice at the time of injection.(*69*) For transfer of Treg cells, Treg cells were sorted from spleen and mLN based on live, CD4^+^FOXP3^+^ROSA^tdTomato+^ expression following pre-enrichment for CD4^+^ cells using negative selection as above. *Rag1^-/-^* mice were intraperitoneally injected with 1x10^5^ Treg cells. For both experiments, mice that received *Trsp*-deficient or -sufficient Tnaïve/Treg cells were co-housed. Mice were weighed at the beginning of each experiment to establish a baseline weight, then weekly thereafter.

#### *In vitro* Treg cell suppression

Naïve T cells (Live CD4^+^CD45RB^hi^) from spleen and mLN were sorted and labeled with CellTrace Violet (ThermoFisher Scientific, #C34557). Treg cells (live CD4^+^CD25^+^ in *CD2^Cre^* mice and live CD4^+^FOXP3^+^ROSA^tdTomato+^ in *Foxp3^eGFP-Cre-ERT2^*^;^ *ROSA26^lox-stop-lox-tdTomato^* mice) were sorted. Cells were stimulated using αCD3αCD28 Dynabeads (ThermoFisher Scientific, #11456D) for approximately 60 hours. Proliferation was assessed via CellTrace Violet dilution in naïve T cells and % suppression determined.

#### Histology

Hematoxylin and eosin (H&E) staining was performed by the VUMC Translational Pathology Shared Resource on formalin-fixed, paraffin-embedded (FFPE) sections. Histological scoring was performed by a gastrointestinal pathologist (MKW) blinded to genotype and treatment.

#### Slide imaging

H&E slides were imaged by the VUMC Digital Histology Shared Resource using a high-throughput Leica SCN400 Slide Scanner automated digital image system from Leica Microsystems. Whole slides were imaged at 40X magnification to a resolution of 0.25 µm/pixel. Images were exported from QuPath(*70*) (version 0.4.3, developed at University of Edinburgh, Edinburgh, Scotland) and if necessary, adjusted for brightness and contrast using Adobe Photoshop CS6 (equally across all images in one figure panel).

#### Mixed bone marrow chimeras

*Rag1^-/-^* recipient mice were pre-conditioned with 450 rads of 137^Cs^ source radiation. Conditioned mice were retro-orbitally injected with 5x10^5^ congenically-marked lineage-depleted (ThermoFisher Scientific, #8804-6829-74) bone marrow cells recovered from the tibiae and femora of CD45.1^+^;*Foxp3^YFP-Cre^* and CD45.2^+^;*Trsp^fl/fl^*;*Foxp3^YFP-Cre^*mice in a 1:1 ratio. Similarly, control mice received cells from CD45.1^+^;*Foxp3^YFP-Cre^*and CD45.2^+^;*Foxp3^YFP-Cre^* mice. All recipient mice received 0.4 mgmL^-1^ enrofloxacin (Baytril, Elanco) in their drinking water from 7 days pre-to 7 days after transplant (14 total days), and 1 mgL^-1^ neomycin from day 3 to day 10 after transplant. Mice were euthanized 6 weeks post-transplant for evaluation.

#### Dextran sodium sulfate colitis

To induce colitis, mice received 3.5% DSS (TdB consultancy, Uppsala, Sweden) in their drinking water for 5 days, followed by 5 days of regular autoclaved drinking water, after which mice were euthanized. Mice were weighed at the beginning of each experiment to establish a baseline weight, then daily thereafter.

### ELISAs

#### IgE ELISA

Serum IgE levels were measured with a Mouse IgE ELISA Kit (ThermoFisher Scientific, #EMIGHE) according to the manufacturer’s protocol, with the following modifications. For *Foxp3^YFP-Cre^;Trsp* samples, the precoated anti-mIgE 96-well plate included in the kit was used. For *Foxp3^scurfy^* samples, a Nunc MaxiSorp 96-well plate (ThermoFisher Scientific, #445101) was coated with 1 µgmL^-1^ anti-mIgE (BioLegend, #406902) overnight at 4° C. The following day, the plate was washed with PBS five times, then blocked with 75% (v/v) PBS with 0.1% Tween 20 (Millipore Sigma, #P9416), 25% (v/v) Block ACE (Bio-Rad, #BUF029), 5% (w/v) α-lactose monohydrate (Millipore Sigma, #L3625), and 5% (w/v) D-sucrose (ThermoFisher Scientific, #BP220-212) for 1 hour at room temperature. All serum samples were diluted 1:10000 in the 1X Assay Diluent included in the kit prior to ELISA. Absorbance at 450 nm was read on a GloMax® Discover plate reader (Promega, Madison, WI).

#### IL-2 ELISA

Samples were diluted (1:15-1:50) and used in the ELISA MAX^TM^ Deluxe Set Mouse IL-2 (Biolegend, #431004) according to manufacturer’s instructions. Absorbance at 450 nm was read on a GloMax® Discover plate reader (Promega).

#### Phosphoprotein assay

CD4^+^ T cells were enriched from the spleen and mLN of tamoxifen-treated mice using negative magnetic selection. Cells were stimulated using 2μgmL^-1^ plate-bound αCD3 (ThermoFisher Scientific, #16-0031-86) and 1 μgmL^-1^ soluble αCD28 (ThermoFisher Scientific, #16-0281-85) for 2 hours. The T-Cell Receptor Phospho Antibody Array (Full Moon Biosystems, #PTC188) was then performed with these cells according to the manufacturer’s protocol, with 10 µLmL^-1^ each of phosphatase inhibitor cocktail 2 (Millipore Sigma, #P5726), phosphatase inhibitor cocktail 3 (Millipore Sigma, #P0044), and protease inhibitor cocktail (Millipore Sigma, #P8340) added to the provided Extraction Buffer for cell lysis. Cy3-streptavidin (ThermoFisher Scientific, #SA1010) was used as the detection antibody. The stained arrays were then imaged, quantified, and analyzed by Full Moon Biosystems. Resulting data displayed is the ratio analysis (ratio change, see description that follows) executed and described by the manufacturer: “For each spot on the array, Median Signal Intensity is extracted from array image. *Average Signal Intensity of Replicate Spots* is the mean value of Median Signal Intensity of the replicate spots for each antibody. Using the *Average Signal Intensity of Replicate Spots*, for each pair of site-specific antibody and phospho site-specific antibody, *Signal Ratio* of the paired antibodies is determined. Ratio = (Signal Intensity of Phospho Site-Specific Antibody) / (Signal Intensity of Site-Specific Antibody). Ratio change between samples = Treatment Sample / Control Sample.”

#### 2-HOBA treatment

*Foxp3^YFP-Cre^;Trsp* mice were provided 1.33 gL^-1^ 2-hydroxybenzylamine acetate (TSI Group Co., Ltd) in the drinking water *ad libitum* from birth.

#### Protein extraction

After anti-CD3/28 stimulation, CD4^+^ T cells were collected and centrifuged at 300 *g* for 5 minutes at 4° C. Cells were resuspended in RIPA Lysis and Extraction Buffer (ThermoFisher Scientific, #89900) with 10 µLmL^-1^ each of phosphatase inhibitor cocktail 2 (Millipore Sigma, #P5726), phosphatase inhibitor cocktail 3 (Millipore Sigma, #P0044), and protease inhibitor cocktail (Millipore Sigma, #P8340), incubated on ice for 10 minutes, and centrifuged at 16000 *g* for 10 minutes at 4° C. Supernatant protein concentrations were quantified with a BCA Protein Assay Kit (ThermoFisher Scientific, #23225) per manufacturer’s instructions.

#### WES^TM^ Simple Western Analysis

Cell lysates at 0.5 µgµl^-1^ were denatured at 95°C for 5 minutes and target proteins detected using an automated capillary-based immunodetection system (WES^TM^ SimpleWestern, Biotechne, Minneapolis, MN) according to the manufacturer’s protocol (ProteinSimple, San Jose, CA). Primary antibodies against GPX4 (abcam, #ab125066), TXNRD1 (abcam, #ab124954) and β-actin (Millipore Sigma, #A5441) were diluted 1:25 in kit-supplied dilution buffer. Wes^TM^ reagents (biotinylated molecular weight marker, streptavidin–HRP fluorescent standards, luminol-S, hydrogen peroxide, sample buffer, DTT, running buffer, wash buffer, matrix removal buffer, secondary antibodies, antibody diluent, and capillaries) were obtained from the manufacturer and used according to the manufacturer’s instructions in supplied partially pre-filled microplates. “Virtual blots” electrophoretic images were automatically generated by the Compass Software and data analysis was performed using the same software (ProteinSimple).

#### TCR activation Western blot

Lysates were diluted in 4X Laemmli Sample Buffer (Bio-Rad, #1610747) with 6% (v/v) 2-mercaptoethanol (Millipore Sigma, #M3148), then incubated at 95° C for 5 minutes. 30 µg total protein was loaded into each lane of a 10-well 4-20% Mini-PROTEAN TGX Precast Protein Gel (Bio-Rad, #4561094), alongside Precision Plus Protein Dual Color Standards (Bio-Rad, #1610374) for SDS-PAGE. Separated proteins were then transferred to a 0.45 µm nitrocellulose membrane (ThermoFisher Scientific, #LC2001), blocked with Intercept (TBS) Blocking Buffer (LI-COR, #927-60001) at room temperature for 30 minutes, then probed with primary antibodies diluted in 50% Intercept (TBS) Blocking Buffer/50% TBS with 0.1% (v/v) Tween-20 (Millipore Sigma, P9416) (TBS-T) at 4°C overnight. Primary antibodies used included: rabbit anti-LCK (1:1000, Cell Signaling Technology, #2984), rabbit anti-pLCK (pTyr394) (1:1000, abcam, #ab318960), mouse anti-MEK1 (1:2000, Cell Signaling Technology, #2352), rabbit anti-pMEK1 (pSer298) (1:1000, Cell Signaling Technology, #9128), and as loading control rabbit anti-vinculin (1:1000, Cell Signaling Technology, #13901). Membranes were washed with TBS-T, then probed with IRDye 680LT goat anti-mouse IgG (1:10,000, LI-COR, #92668020) and IRDye 800CW goat anti-rabbit IgG (1:10,000, LI-COR, #92632211) secondary antibodies diluted in TBS-T at room temperature for 30 minutes. Membranes were washed again with TBS-T, imaged with an Odyssey Clx near-infrared fluorescence imaging system (LI-COR), and quantified with Image Studio (LI-COR). Densitometric values for proteins of interest were normalized to those of their corresponding loading controls.

#### Treg cell survival assay

Treg cells (Live CD4^+^FOXP3^+^ROSA^tdTomato+^) were sorted from the spleen and mLN of tamoxifen-treated mice. 4x10^4^ cells were plated in each well of a round-bottom 96-well plate with human IL-2 (Biolegend or R&D Systems) at concentrations indicated in the figure legend. After 48 hours, CellTiter-Glo® Luminescent Cell Viability Assay (Promega, #G7572) was performed according to manufacturer’s instruction, and results normalized to TCM only negative control.

#### Human thymus single cell RNA-seq analysis

A public single-cell RNA-seq dataset of human thymocytes(*31*) was manually transcribed from the web-based interface to Prism (Version 10.1.1).

#### TCRβ chain immunosequencing

Samples were enriched for CD4^+^ T cells using negative selection as above. Cells were then stained and flow-sorted into PBS with 2% FBS and 0.025 M HEPES followed by DNA extraction using the DNeasy Blood and Tissue kit (Qiagen, Hilden, Germany, #69504). TCR beta sequencing was performed by Adaptive Biotech using the immunoSEQ Assay. Raw data was analyzed using the immunoSEQ Analyzer 3.0 website (clients.adaptivebiotech.com/login).

#### RNA isolation, cDNA synthesis, and RT-qPCR

RNA extraction from cells was performed using the RNeasy Mini Kit (Qiagen, #4304437). For low cell numbers the RNAeasy Micro Kit (Qiagen, #74004) was used, and cells were sorted directly into 300 μL RLT buffer (Qiagen, #79216) containing 2-mercapto-ethanol as recommended. On column DNAse treatment was performed as per manufacturer’s recommendations. For cDNA synthesis, 5 µL RNA, 4 µL SuperScript IV VILO Master Mix (ThermoFisher Scientific, #11756050), and 11 µL water were combined in a PCR tube and incubated as follows: 1) at 25° C for 10 minutes, 2) at 50° C for 10 minutes, and 3) at 85° C for 5 minutes. Selenoprotein RT-qPCR was performed in technical duplicate using TaqMan™ probes described previously by our group(*30*) and TaqMan™ Universal PCR Master Mix (ThermoFisher Scientific, #4304437) on a QuantStudio3 (ThermoFisher Scientific). Selenoprotein RT-qPCR results were analyzed by the delta-delta Ct method and normalized to *Tbp*. For *Lef1* RT-qPCR, SYBR primers (Millipore Sigma) (5’ TGTTTATCCCATCACGGGTGG 3’ and 5’ CATGGAAGTGTCGCCTGACAG 3’) and PerfeCTa® SYBR® Green SuperMix ROX (Quantabio, #9505502K) were used. *Lef1* RT-qPCR results were normalized to *Ubc* (5’ GCCCAGTGTTACCACCAAGA 3’ and 5’ CCCATCACACCCAAGAACA 3’).

#### RNA isolation for sequencing

Quality control for RNA was performed by the Vanderbilt Technologies for Advanced Genomics core using RNA 6000 Pico (Agilent, Santa Clara, CA), and cDNA library preparation using a NEB library preparation kit. Paired end 150bp sequencing was performed on a NovaSeq 6000 (Illumina).

#### RNA-sequencing analysis (bulk)

Briefly, samples were trimmed with fastp (version 0.20.0) using default parameters. Quantification was performed using Salmon (version 1.4.0) against a decoy transcriptome (Mus musculus Gencode version 21). Further analysis was performed in R (version 4.1.2) in R studio (version 2021.09.2 + 382). For differential expression analysis, limma (version 3.50.3) was used on log-CPM transformed counts with prior count set to 3 or DESEq2 (version 1.34.0) was used on non-normalized counts.

#### T cell polarization

T cells were acquired from spleen and mLN of WT C57/Bl6 mice using negative enrichment. T cells were then activated using 2μgmL^-1^ plate-bound αCD3 (ThermoFisher Scientific, #16-0031-86) and 1 μgmL^-1^ soluble αCD28 (ThermoFisher Scientific, #16-0281-85) and the following cytokines: Th0: 20 ngmL^-1^ IL-2 (Biolegend, #575406). Th1: 20 ngmL^-1^ IL-2, 10 ngmL^-1^ IL-12 (R&D Systems, #419-ML-010/CF), and 2 μgmL^-1^ αIL-4 (ThermoFisher Scientific, #16-7041-85). Th2: 20 ngmL^-1^ IL-2, 10 ngmL^-1^ IL-4 (R&D Systems, #404-ML-010/CF), and 1 μgmL^-1^ αIFNγ (ThermoFisher Scientific, #14-7311-85). Treg: 40 ngmL^-1^ IL-2, 1 μgmL^-1^ αIFNγ, 1 μgmL^-1^ αIL-4, and 2 ngmL^-1^ human TGFβ (R&D Systems, #240-B-002/CF). Th17: 20 ngmL^-1^ IL-6 (R&D Systems, #406-ML-005/CF), 1 ngmL^-1^ human TGFβ, 1 μgmL^-1^ αIL-12 (BD, #554475), 1 μgmL^-1^ αIFNγ, and 1 μgmL^-1^ αIL-4. Th0, Th1, Th2, and Treg cells were cultured in RPMI 1640, while Th17 cells were cultured in IMDM (ThermoFisher Scientific, #12440079) base media. All T cell media contained 10% (v/v) heat-inactivated FBS, 100 unitsmL^-1^ penicillin, 100 μgmL^-1^ streptomycin, 1% (v/v) GlutaMax, 1% (v/v) MEM non-essential amino acids, 55 μM beta-mercaptoethanol, and 1 mM sodium pyruvate.

#### Quantification and statistical analysis

Age-, sex- and, when feasible, littermate-matched mice were used for all experiments. Statistical analysis was performed using GraphPad Prism (9.1.2) unless otherwise described. Sample size was determined empirically. All data points are biological replicates, i.e. each datapoint represents one mouse, unless otherwise indicated, i.e. data from multiple mice were pooled. All figures represent similar results from at least three independent experiments unless otherwise indicated.

## Supplementary Materials

Figs S01-Figure S09

Table S1

## Abbreviations

DSS: dextran sodium sulphate
DN: double negative (CD4^neg^, CD8^neg^)
DP: double positive (CD4^+^, CD8^+^)
FC: flow cytometry
mLN: mesenteric lymph node
RT-qPCR: reverse transcription quantitative polymerase chain reaction
ROS: reactive oxygen species
SI: small intestine
SP: single positive
TCR: T cell receptor
Th: T helper
WT: wild-type

## Acknowledgements

Luc van Kaer, Danyvid Olivares-Villagomez, Kristen Hoek, Kathleen McClanaham for scientific discussions. The Nashville VA Medical Center Flow Core (Christian Warren) and the VUMC Flow Core. VUMC Division of Animal Care, the Translational Pathology Shared Resource, and the Advance Computing Center for Research and Education. Mitanshu (Mit) Pandya for assistance with mouse management. Chase Jones, Nathaniel Berle, Roberta Norris for general lab management. Sophia Ballinger for administrative assistance. The Digital Histology Shared Resource for slide scanning. John Rathmacher from MTI BioTech Inc for provision of 2-HOBA.

## Funding

VA-Career Development Award IK2BX004648 to YAC

Vanderbilt Digestive Diseases Research Center through DDRC Pilot and Feasibility Grant and Complementary

Award P30 058404 to YAC

Crohn’s and Colitis Foundation SRA 1061046 to JAG

NIH NAIAD R01 AI176521 to JAG

VA MERIT Award 2I01BX001426 to CSW

NIH NIDDK R01 DK131810 to CSW

NCI grant P30 CA068485 to CSW

NIDDK T32DK007673 to MAB

Vanderbilt Burroughs Wellcome Fund Supporting Careers in Research for Interventional Physicians and Surgeons Award to MAB

Eunice Kennedy Shriver National Institute of Child Health & Human Development of the National Institutes of Health under Award Number K12HD087023 to MAB

## Author contributions

Conceptualization: JJ and YAC

Data curation: JJ

Formal analysis: JJ, SS, MKW, JMP

Funding acquisition: JAG and YAC

Investigation: JJ, JMP, ABH, DN, SPS, MAB, WD

Methodology: JJ, JMP, JNS, JAG, YAC

Mouse management: AK, ZA, MAB, JMP, JJ, YAC

Project administration: JJ, JMP

Resources: CRF, WD, JCR, KSP, JAG, YAC

Software: JJ, CC, SS

Supervision: JAG, JNS, CSW, YAC

Validation: JJ, JMP, SPS, MAB

Visualization: JJ

Writing – original draft: JJ

Writing – editing: JMP, JAG, YAC

Writing – review: all authors

## Data and materials availability

Bulk RNA-sequencing data generated as part of this study is deposited in GEO: GSE275428 reviewer password: gpivwoaupzspnyv. Code will be deposited at GitHub prior to publication. Other materials are available upon request from the corresponding author.

## Supplementary Materials

**Figure S01.**
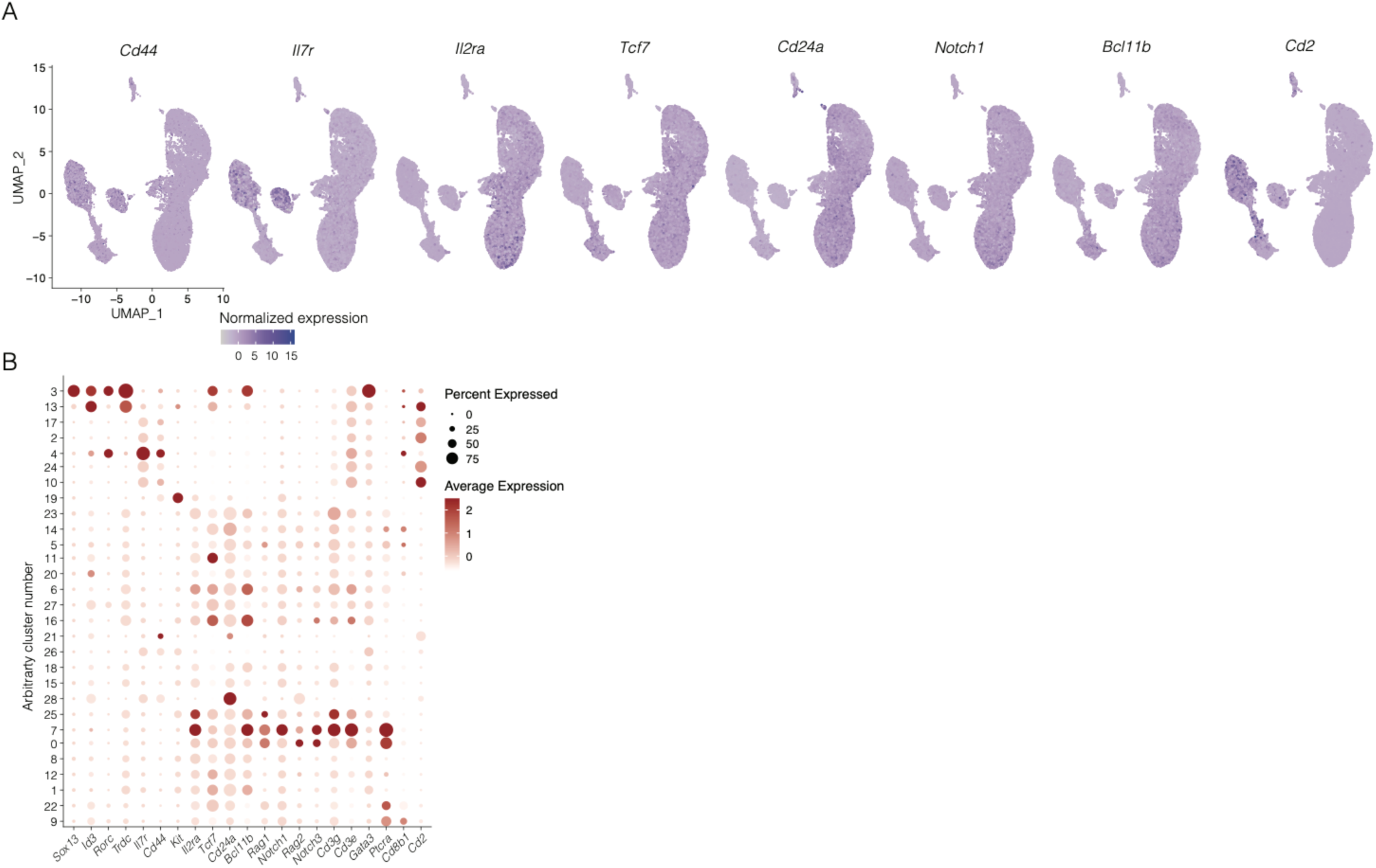
*Cd2* is expressed by mouse DN1 thymocytes. A previously published single cell RNA-seq dataset of mouse DN thymocytes was analyzed similarly as described (see methods for details).^27^ (A) UMAP of clustered cells. DN1 cells were defined as expressing high *Cd44* and *Il7r* and low *Il2ra*, *Tcf7*, *Cd24*, *Notch1,* and *Bcl11b*. (B) Dot plot showing genes used to define clusters of cells representing DN1 thymocytes, i.e. cluster 2, 4, 10, 7, 24 in this analysis. Note that assigned cluster numbers are arbitrary and do not correspond to cluster numbers in the original manuscript.^27^

**Figure S02.**
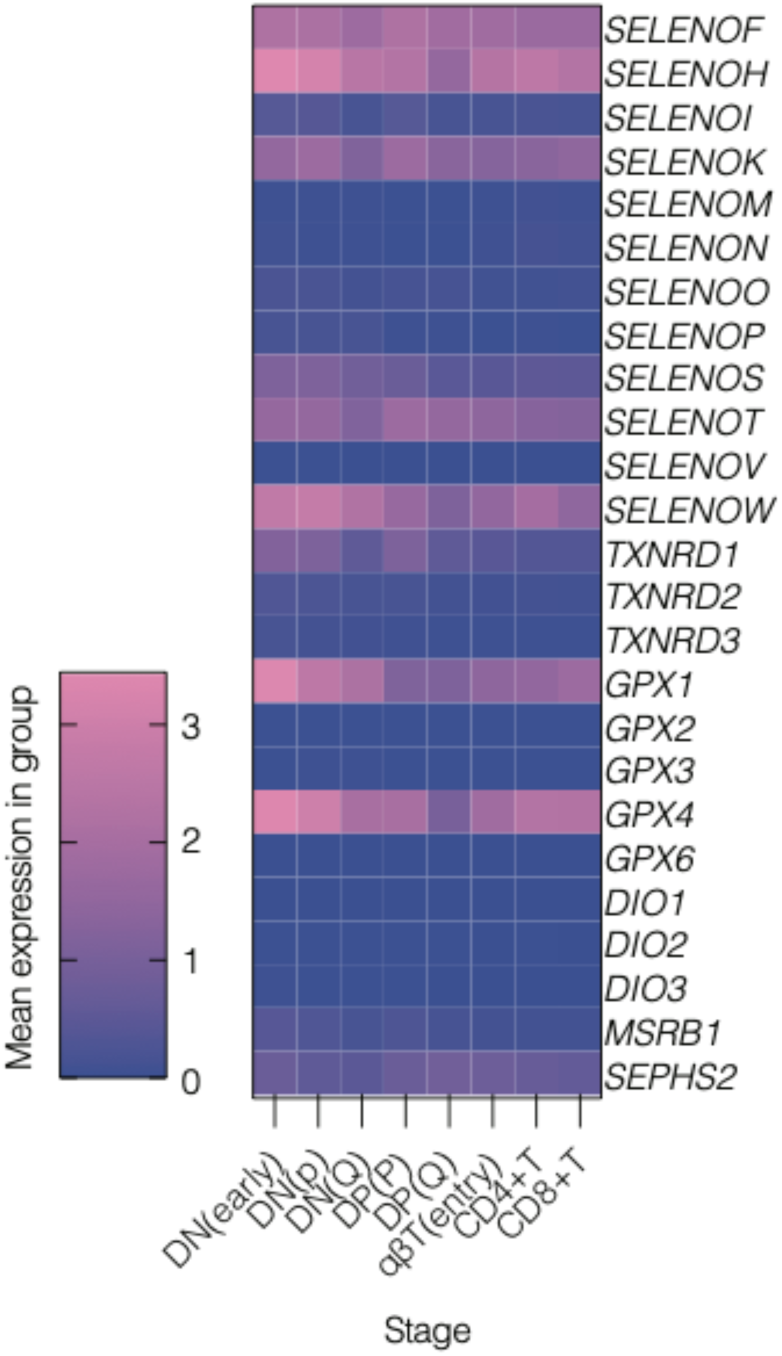
Selenoproteins are expressed by human thymocytes. A previously published single-cell RNA-seq dataset of human thymocytes was analyzed for selenoprotein expression using the same clusters as in the original manuscript^31^.

**Figure S03.**
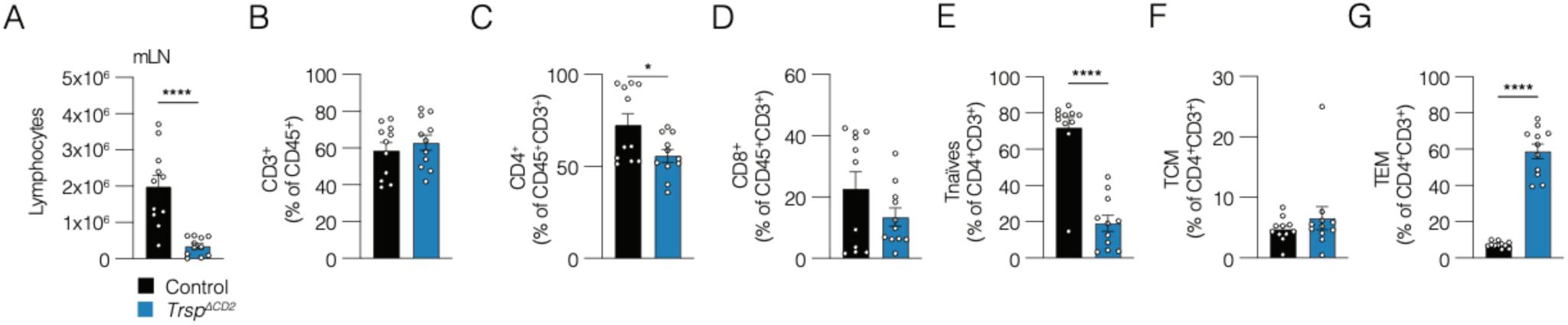
CD2^+^ cells require *Trsp* for T cell homeostasis. (A) Absolute cell count of live lymphocytes in mLN from control and *Trsp^1′CD2^* mice acquired using automatic cell counter. Pooled data from 2 independent experiments. Unpaired t-test. (B-G) FC of mLN lymphocytes from control and *Trsp^1′CD2^* mice. Unpaired t tests. Gating similar to gating described in Figure 1. Pooled data from 2 independent experiments. * p<0.05; ** p<0.01; *** p<0.001; **** p<0.0001.

**Figure S04.**
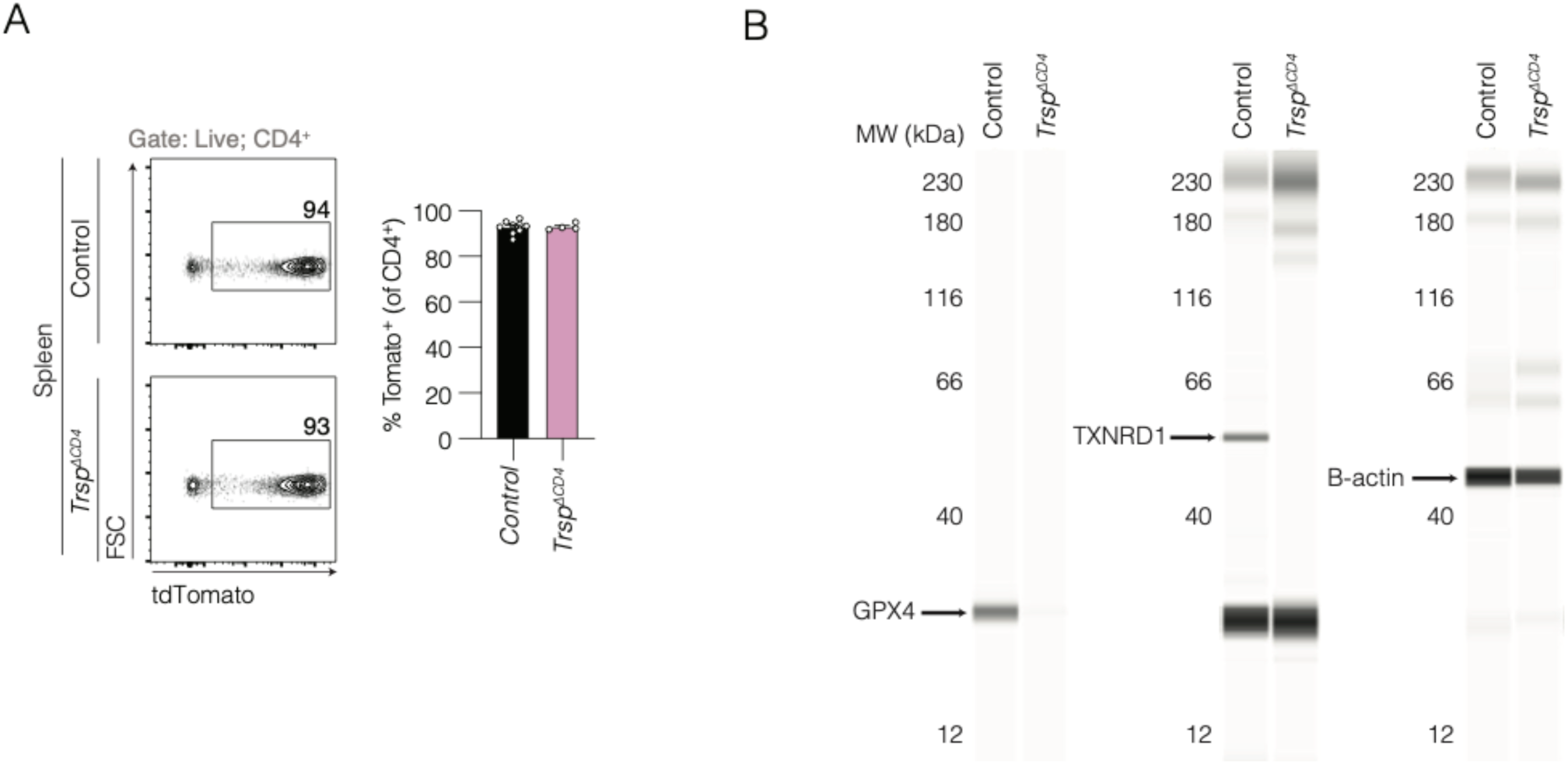
Validation of selenoprotein deletion in *Trsp^1′CD4^* CD4^+^ T cells. (A) Representative FC plots (left) and quantification (right) of T cells from control and *Trsp^ΔCD4^* mice for tdTomato. Data representative of >10 independent experiments. Unpaired t-test. (B) Capillary electrophoresis immunoassay (“Simple Western”) of control and *Trsp^ΔCD4^*CD4^+^ T cells from spleen and mLN acquired using immunomagnetic negative selection for GPX4 (∼22 kDa) or TXNRD1 (∼55 kDa). As this automated system transfers an equal volume of the same sample into each capillary, B-actin (∼42 kDa) was used as a loading control in separate capillaries. Samples were pooled from n=4-5 mice per genotype, and data are representative from 2 independent experiments.

**Figure S05.**
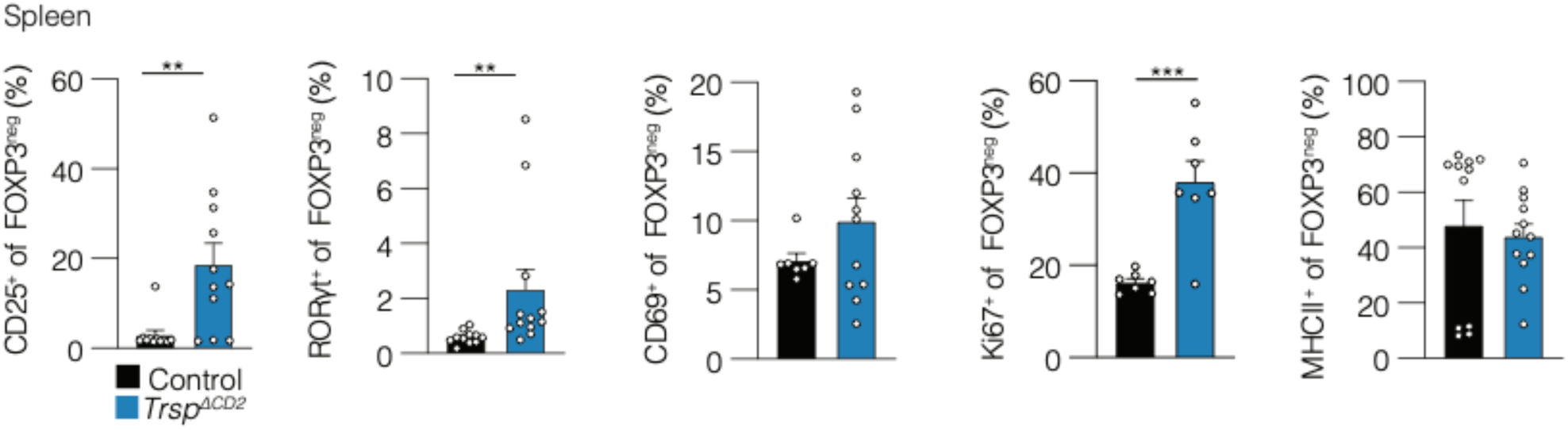
Phenotyping of FOXP3^neg^ CD4^+^ T cells in *Trsp^1′CD2^* mice. FC quantification of splenocytes from control and *Trsp^1′CD2^* mice from Figure 3 with similar gating, showing frequencies of FOXP3^neg^CD4^+^ T cells. Unpaired t tests. **p<0.01, ***p<0.001.

**Figure S06.**
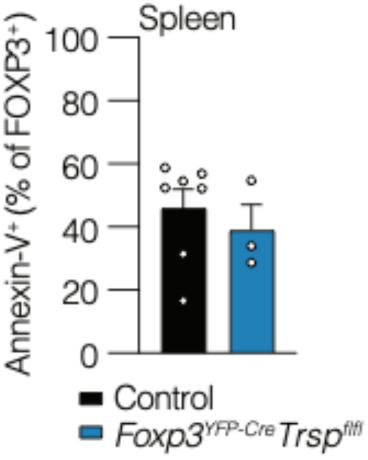
Phenotyping of *Trsp*-deficient Treg cells. FC quantification of Annexin-V^+^ Treg cells from control and *Trsp^fl/fl^Foxp3^YFP-Cre^* mice. Mann-Whitney U test.

**Figure S07.**
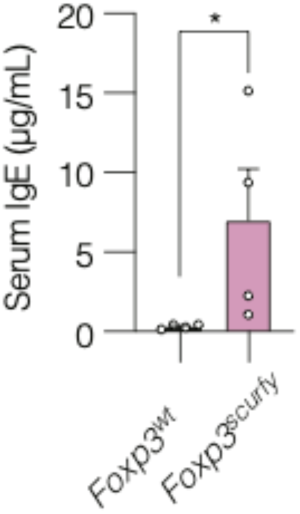
Autoimmune inflammation in *Foxp3^Scurfy^* mice. Serum IgE ELISA in control and *Foxp3^scurfy^* mice. Mann-Whitney U test.

**Figure S08.**
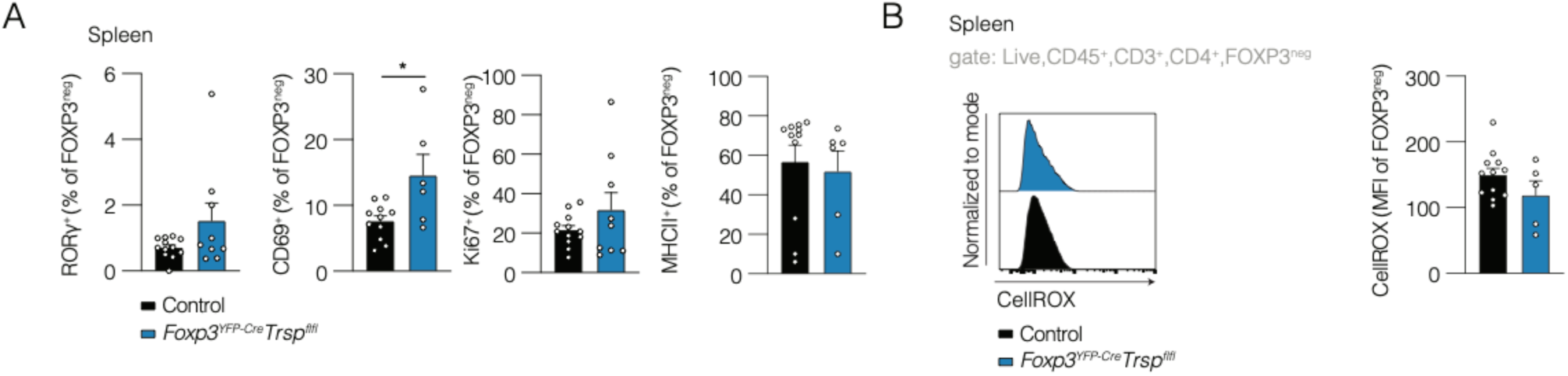
Phenotyping of FOXP3^neg^ CD4^+^ T cells in *Trsp^fl/fl^Foxp3^YFP-Cre^*mice. (A) FC quantification of splenocytes from control and *Trsp^fl/fl^Foxp3^YFP-Cre^*mice from Figure 3 with similar gating, showing frequencies of FOXP3^neg^CD4^+^ T cells. (B) Representative FC histograms (left) and quantification (right) of CellROX fluorescence of FOXP3^neg^CD4^+^ T cells from control and *Trsp^fl/fl^Foxp3^YFP-Cre^*mice. Unpaired t tests. *p<0.05.

**Figure S09.**
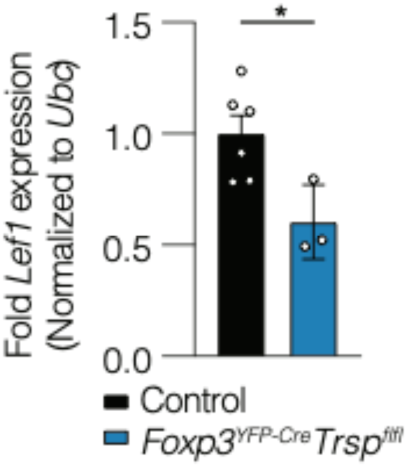
*Lef1* expression in *Trsp*-deficient Treg cells. FOXP3^+^ Treg cells from *Trsp^fl/fl^Foxp3^YFP-Cre^* and control mice were flow-sorted before RT-qPCR for *Lef1* normalized to *Ubc*. Unpaired t-test. One experiment.

## Supplementary table

**Table S1.**
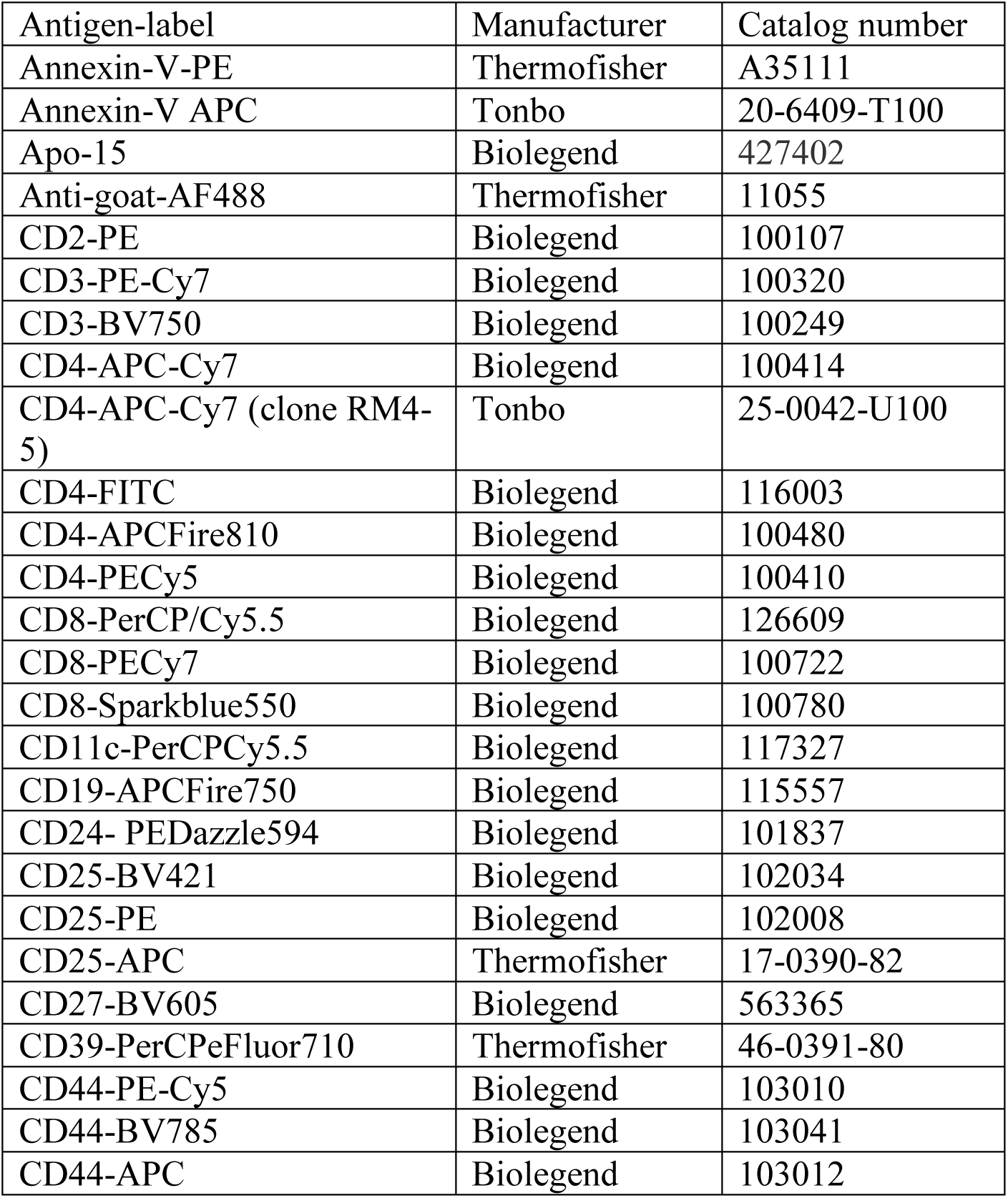
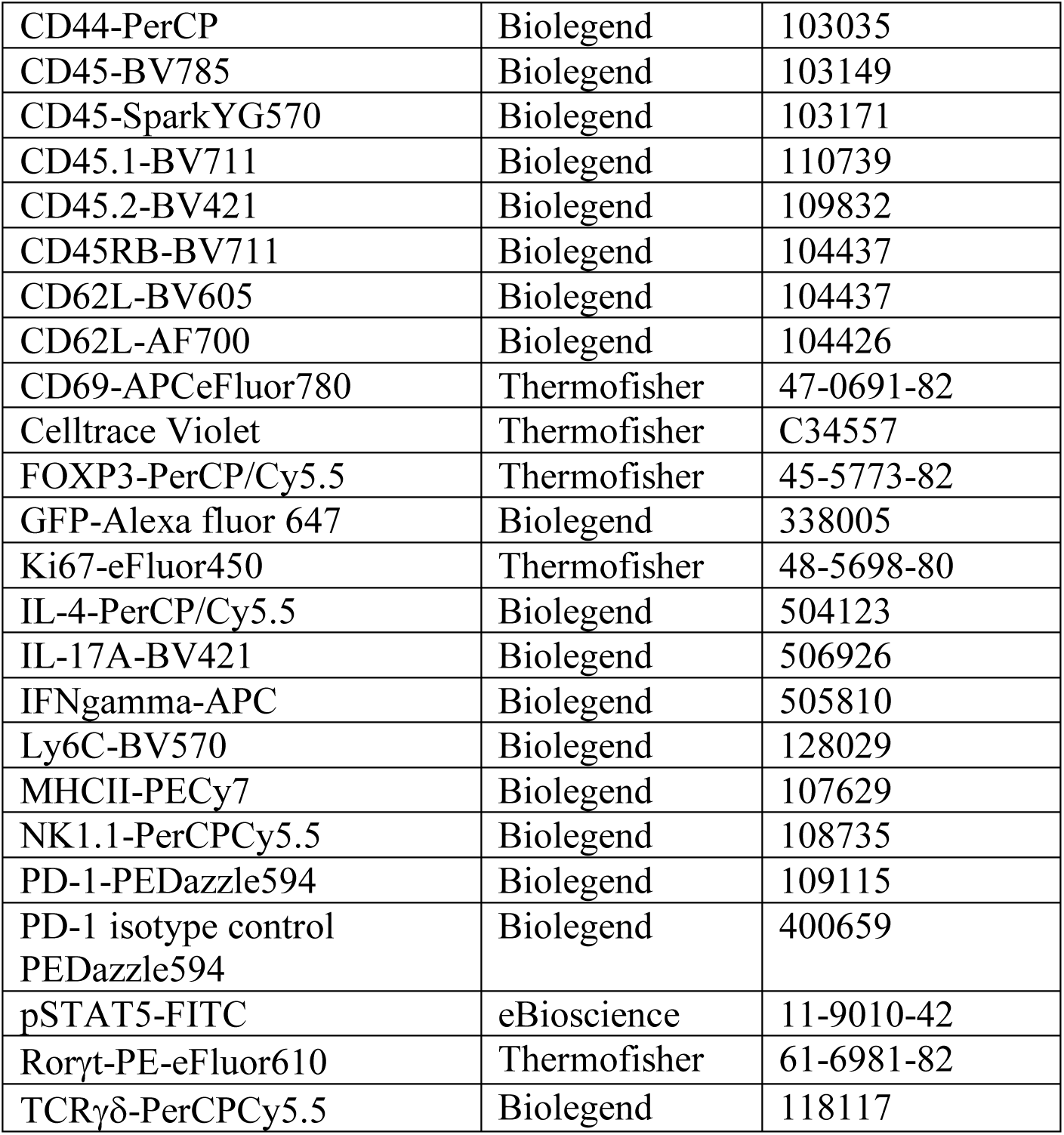
FC antibodies.

