## Supplementary material for "*Trsp* is required by regulatory T cells to prevent lethal autoimmunity in mice": supp files

### Supplementary Materials

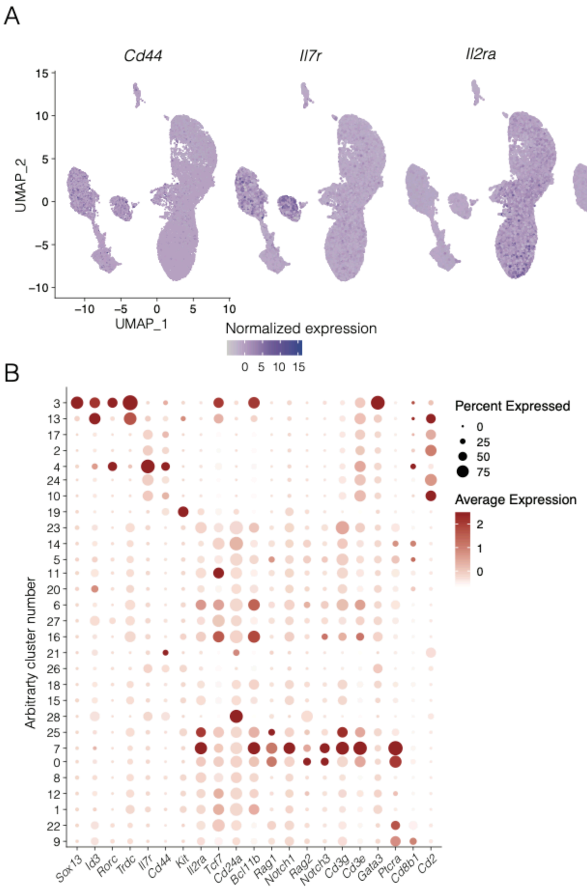


**Figure S01 *Cd2* is expressed by mouse DN1 thymocytes**

A previously published single cell RNA-seq dataset of mouse DN thymocytes was analyzed similarly as described (see methods for details).[27] (A) UMAP of clustered cells. DN1 cells were defined as expressing high *Cd44* and *Il7r* and low *Il2ra*, *Tcf7*, *Cd24*, *Notch1,* and *Bcl11b*.

(B) Dot plot showing genes used to define clusters of cells representing DN1 thymocytes, i.e. cluster 2, 4, 10, 7, 24 in this analysis. Note that assigned cluster numbers are arbitrary and do not correspond to cluster numbers in the original manuscript.[27]

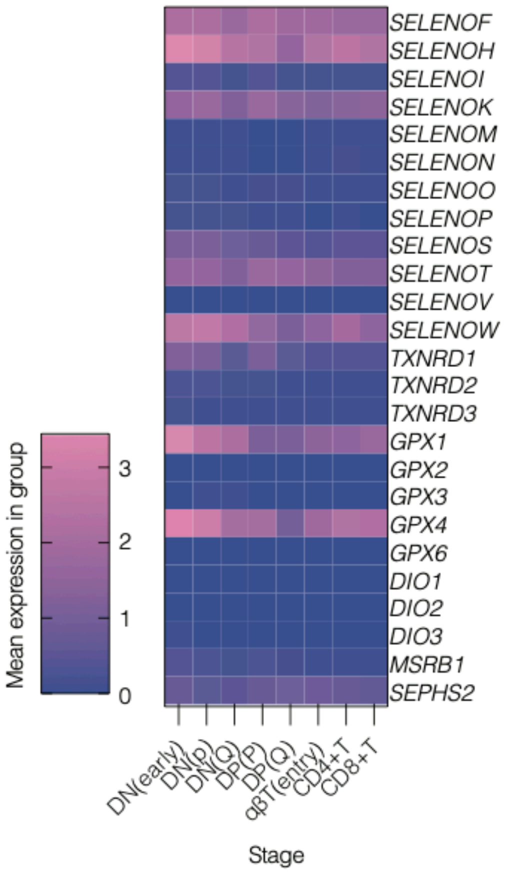


**Figure S02 Selenoproteins are expressed by human thymocytes.**

A previously published single-cell RNA-seq dataset of human thymocytes was analyzed for selenoprotein expression using the same clusters as in the original manuscript[31].

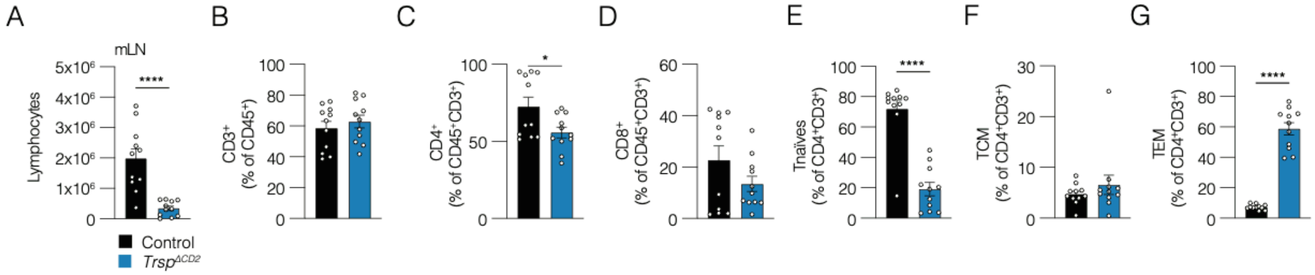


**Figure S03 CD2+ cells require *Trsp* for T cell homeostasis.**

(A) Absolute cell count of live lymphocytes in mLN from control and *Trsp*$^{\Delta CD2}$ mice acquired using automatic cell counter. Pooled data from 2 independent experiments. Unpaired t-test.

(B-G) FC of mLN lymphocytes from control and *Trsp*$^{\Delta CD2}$ mice. Unpaired t tests. Gating similar to gating described in **Figure 1**. Pooled data from 2 independent experiments.

* $p<0.05$; ** $p<0.01$; *** $p<0.001$; **** $p<0.0001$.

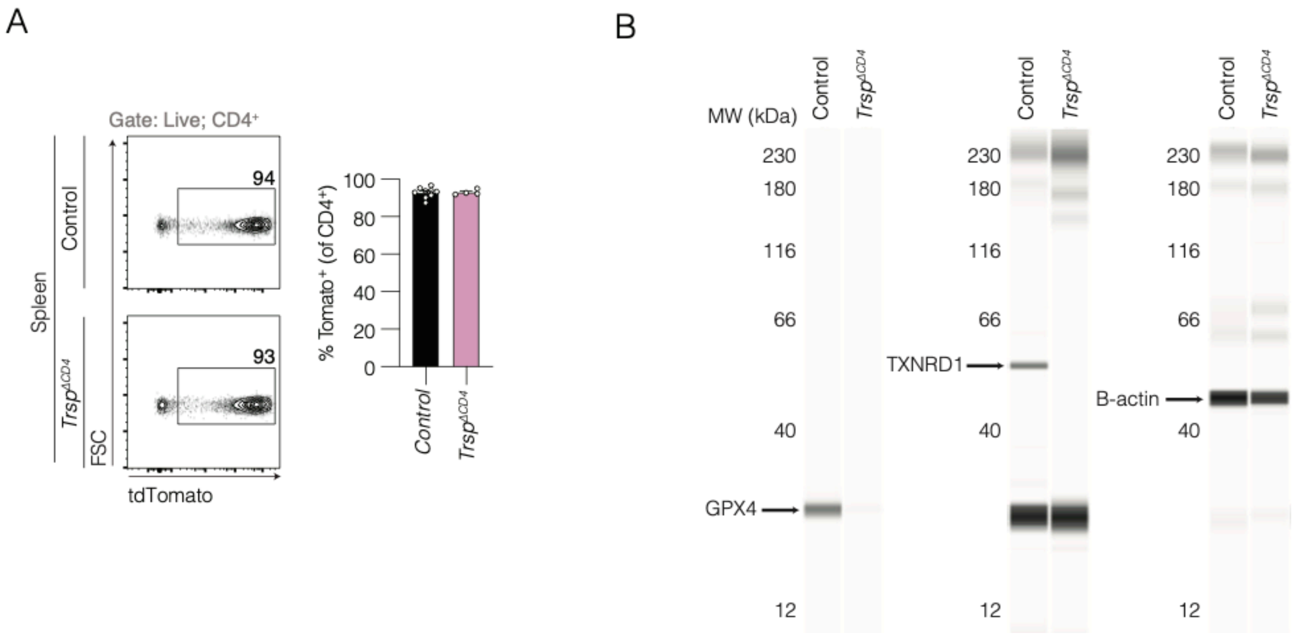


**Figure S04 Validation of selenoprotein deletion in *$Trsp^{\Delta CD4}$* $CD4^+$ T cells**

(A) Representative FC plots (left) and quantification (right) of T cells from control and *$Trsp^{\Delta CD4}$* mice for tdTomato. Data representative of >10 independent experiments. Unpaired t-test.

(B) Capillary electrophoresis immunoassay ("Simple Western") of control and *$Trsp^{\Delta CD4}$* $CD4^+$ T cells from spleen and mLN acquired using immunomagnetic negative selection for GPX4 (~22 kDa) or TXNRD1 (~55 kDa). As this automated system transfers an equal volume of the same sample into each capillary, B-actin (~42 kDa) was used as a loading control in separate capillaries. Samples were pooled from n=4-5 mice per genotype, and data are representative from 2 independent experiments.

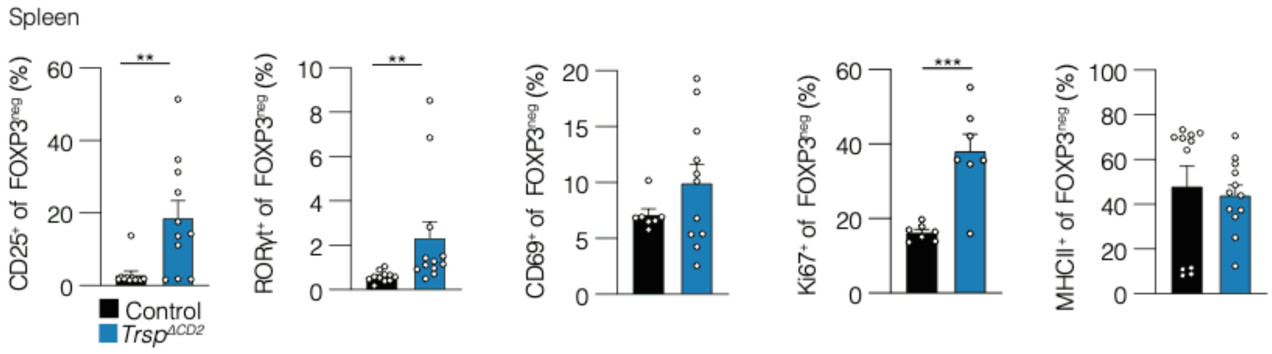


**Figure S05 Phenotyping of FOXP3^neg^ CD4^+^ T cells in *Trsp^ΔCD2^* mice.**

FC quantification of splenocytes from control and *Trsp^ΔCD2^* mice from **Figure 3** with similar gating, showing frequencies of FOXP3^neg^CD4^+^ T cells. Unpaired t tests. **p<0.01, ***p<0.001.

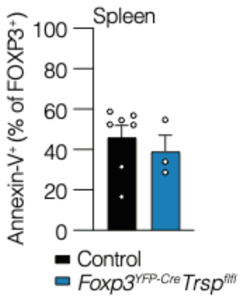


**Figure S06 Phenotyping of *Trsp*-deficient Treg cells.**

FC quantification of Annexin-V$^{+}$ Treg cells from control and $Trsp^{fl/fl}Foxp3^{YFP\text{-}Cre}$ mice. Mann-Whitney U test.

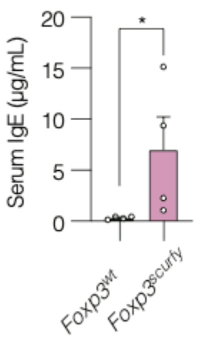


**Figure S07 Autoimmune inflammation in *Foxp3<sup>Scurfy</sup>* mice**

Serum IgE ELISA in control and *Foxp3*$^{scurfy}$ mice. Mann-Whitney U test.

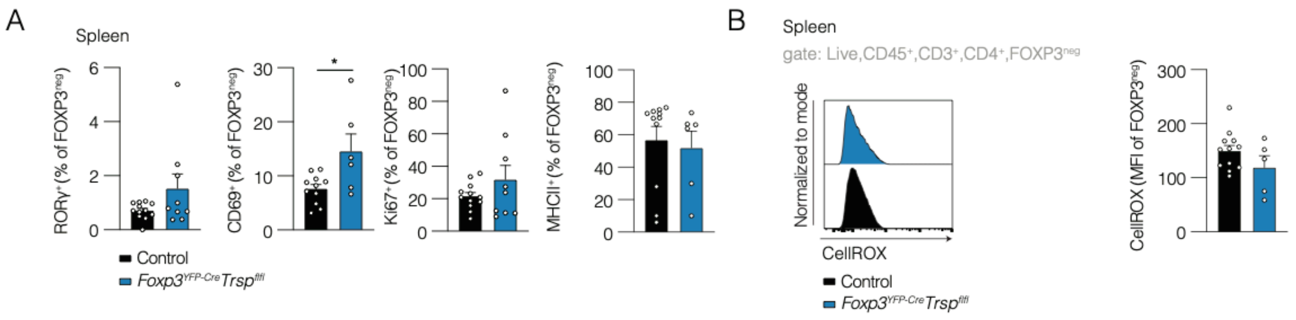


**Figure S08 Phenotyping of FOXP3$^{neg}$ CD4$^{+}$ T cells in *Trsp$^{fl/fl}$Foxp3$^{YFP\text{-}Cre}$* mice**

(A) FC quantification of splenocytes from control and *Trsp$^{fl/fl}$Foxp3$^{YFP\text{-}Cre}$* mice from Figure 3 with similar gating, showing frequencies of FOXP3$^{neg}$CD4$^{+}$ T cells. (B) Representative FC histograms (left) and quantification (right) of CellROX fluorescence of FOXP3$^{neg}$CD4$^{+}$ T cells from control and *Trsp$^{fl/fl}$Foxp3$^{YFP\text{-}Cre}$* mice. Unpaired t tests. *p<0.05.

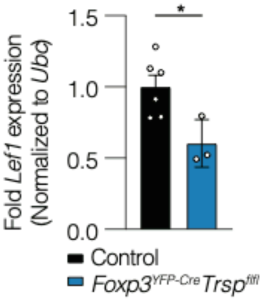


**Figure S09** ***Lef1*** **expression in** ***Trsp*****-deficient Treg cells**

FOXP3$^{+}$ Treg cells from *Trsp*$^{fl/fl}$*Foxp3*$^{YFP\text{-}Cre}$ and control mice were flow-sorted before RT-qPCR for *Lef1* normalized to *Ubc*. Unpaired t-test. One experiment.

### Supplementary table

**Table S1 FC antibodies**

| Antigen-label | Manufacturer | Catalog number |
|---|---|---|
| Annexin-V-PE | Thermofisher | A35111 |
| Annexin-V APC | Tonbo | 20-6409-T100 |
| Apo-15 | Biolegend | 427402 |
| Anti-goat-AF488 | Thermofisher | 11055 |
| CD2-PE | Biolegend | 100107 |
| CD3-PE-Cy7 | Biolegend | 100320 |
| CD3-BV750 | Biolegend | 100249 |
| CD4-APC-Cy7 | Biolegend | 100414 |
| CD4-APC-Cy7 (clone RM4-5) | Tonbo | 25-0042-U100 |
| CD4-FITC | Biolegend | 116003 |
| CD4-APCFire810 | Biolegend | 100480 |
| CD4-PECy5 | Biolegend | 100410 |
| CD8-PerCP/Cy5.5 | Biolegend | 126609 |
| CD8-PECy7 | Biolegend | 100722 |
| CD8-Sparkblue550 | Biolegend | 100780 |
| CD11c-PerCPCy5.5 | Biolegend | 117327 |
| CD19-APCFire750 | Biolegend | 115557 |
| CD24- PEDazzle594 | Biolegend | 101837 |
| CD25-BV421 | Biolegend | 102034 |
| CD25-PE | Biolegend | 102008 |
| CD25-APC | Thermofisher | 17-0390-82 |
| CD27-BV605 | Biolegend | 563365 |
| CD39-PerCPeFluor710 | Thermofisher | 46-0391-80 |
| CD44-PE-Cy5 | Biolegend | 103010 |
| CD44-BV785 | Biolegend | 103041 |
| CD44-APC | Biolegend | 103012 |

| | | |
|---|---|---|
| CD44-PerCP | Biolegend | 103035 |
| CD45-BV785 | Biolegend | 103149 |
| CD45-SparkYG570 | Biolegend | 103171 |
| CD45.1-BV711 | Biolegend | 110739 |
| CD45.2-BV421 | Biolegend | 109832 |
| CD45RB-BV711 | Biolegend | 104437 |
| CD62L-BV605 | Biolegend | 104437 |
| CD62L-AF700 | Biolegend | 104426 |
| CD69-APCeFluor780 | Thermofisher | 47-0691-82 |
| Celltrace Violet | Thermofisher | C34557 |
| FOXP3-PerCP/Cy5.5 | Thermofisher | 45-5773-82 |
| GFP-Alexa fluor 647 | Biolegend | 338005 |
| Ki67-eFluor450 | Thermofisher | 48-5698-80 |
| IL-4-PerCP/Cy5.5 | Biolegend | 504123 |
| IL-17A-BV421 | Biolegend | 506926 |
| IFNgamma-APC | Biolegend | 505810 |
| Ly6C-BV570 | Biolegend | 128029 |
| MHCII-PECy7 | Biolegend | 107629 |
| NK1.1-PerCPCy5.5 | Biolegend | 108735 |
| PD-1-PEDazzle594 | Biolegend | 109115 |
| PD-1 isotype control PEDazzle594 | Biolegend | 400659 |
| pSTAT5-FITC | eBioscience | 11-9010-42 |
| Rorγt-PE-eFluor610 | Thermofisher | 61-6981-82 |
| TCRγδ-PerCPCy5.5 | Biolegend | 118117 |
